# Improved tests for the origin of allometric scaling across tree architectures

**DOI:** 10.1101/2024.07.25.605048

**Authors:** Adam Chmurzynski, Alexander Byers Brummer, Van Savage, Alexander Shenkin, Yadvinder Malhi, Olivier Martin-Ducup, Kasia Zieminska, Nicolas Barbier, Brian J. Enquist

## Abstract

The scaling of organismal metabolic rates with body size is one of the most prominent empirical patterns in biology. For over a century, the nature and causes of metabolic scaling have been the subject of much focus and debate. West, Brown, and Enquist (WBE) proposed a general model for the origin of metabolic scaling from branching vascular networks. However, recent empirical tests of WBE vascular scaling predictions in plants and animals have reported deviations caused by variability in network geometry. After clarifying the core assumptions of the WBE model, we revisit the methods and conclusions of recent tests conducted in trees, finding support for key WBE predictions in woody plant architecture. To do this, we apply an approach that better captures: i) network branching self-similarity and ii) leaf area as a proxy of plant metabolic capacity. The WBE model also predicts curvature in metabolic scaling in smaller organisms, and we introduce a novel method that accounts for curvature in plant branching geometry. Together, these advances allow more direct measurements of metabolic scaling than previous work, and we apply them to a dataset of diverse laser-scanned tree architectures. Analyses reveal the predicted interspecific ¾ metabolic scaling across tree crowns, with intraspecific variation within individual tree crowns. Scaling variability is consistent with WBE predictions for curvature from asymptotic growth and underlying variation in branching geometry. We conclude that linking fine-scale branching variation to metabolic scaling allometries remains a challenge, while our results support the foundational hypotheses of the WBE model.

**Author summary:** Trees survive in a variety of habitats and lifestyles across Earth. They are also characterized by a stunning array of sizes and shapes that make trees objects of vast cultural, economic, and ecological importance. At the same time, the need to link vascular plant function with traits and environment is more pressing than ever. Size (body mass) is fundamentally linked to plant functioning within ecosystems through allometric relationships. Allometric relationships emerge from the geometry of branch networks in trees, which are increasingly well-characterized with remote-sensing data. We use a dataset of laser-scanned tree crowns to test allometric predictions that link size to key traits, particularly metabolic capacity, understood as total leaf area. Our results indicate that i) scanning technology can provide accurate assessments of branch allometry with proper data preparation, and ii) studying branch allometries provides an organizing framework for interpreting natural variation in tree architecture.

## Introduction

During its lifespan, a forest tree can span the greatest range of body sizes of any organism that has ever lived on Earth. From the smallest seedlings weighing less than 1 gram to the tallest canopy trees weighing hundreds of megagrams, an individual tree can span up to 12 orders of magnitude in mass [1]. Moreover, forests are among the largest terrestrial carbon pools on the planet, playing a pivotal role in annual carbon cycling [2]. The diversity of branch and crown geometries has long suggested a link with variation in life history resource strategies of trees [3–5]. Variation in size and shape at scales ranging from leaves to whole crowns (hereafter, tree branching architecture) is thought to play a key role in how trees respond to a variable light and resource environment, influencing growth, competition, and community assembly across forest ecosystems [6–8].

Allometric scaling relationships can capture and explain variation in size and shape among organisms by describing how biological traits scale with body size. Specifically, *Y=Y*_*0*_ *M*^*b*^ where *Y* is the dependent variable, *M* is body mass, *b* is a power exponent, and *Y*_*0*_ is a normalization constant that varies with the nature of *Y* and possibly with the kind of organism.

Such allometric scaling relationships provide a foundation for scaling the physiology of trees up to the dynamics of forests [9–12]. Studies of plants and animals have shown that many biological traits scale with quarter-powers of mass, for example, *b* ∼ 3/4 for metabolic rate and growth rate, with implications for ecology and life history [13–15]. In plants, analyses of gas exchange measures of respiration rates across broad size ranges show a central tendency of ¾ scaling relationships [16]. Several large-scale interspecific comparative studies support the core assertion of quarter-power scaling across broad domains of life [17–20]. Nonetheless, ecological analyses of plants have also shown that broad variation in allometric scaling relationships should be expected from environmental pressures [21–25].

West, Brown, and Enquist (WBE) proposed a general model for the origin of allometric scaling laws in biology [26]. The model assumes a hierarchical network, branching from base to tips, with terminal segments serving matter and energy to cells through vascular transport.

According to WBE, whole-organismal metabolism and growth rate are under stabilizing selection that minimizes the scaling of hydrodynamic resistance and maximizes the scaling of resource uptake [26]. The evolutionary outcome of this optimization is a vascular network that is fractal-like and characterized by a small number of self-similar branching traits that are constrained to specific values. Critically, those branching traits predict that whole-organism metabolic rate scales as the ¾ power of body mass.

WBE’s most intriguing implication is that relaxation of stabilizing selection could permit variation in vascular branching traits, predicting variation in allometric scaling relationships across levels of organization [11,27,28]. WBE predicts that degrees of variation in branching traits cause specific deviations in allometric scaling relationships and in particular deviation from the ¾ scaling of ecologically relevant, dynamic organismal attributes such as photosynthesis, metabolism, and growth rates [29,30]. Depending on the local environment, deviations above and below the ¾ scaling exponent might be advantageous. For example, directional selection for allometric adaptation may be due to selection for faster growth and a short lifespan to escape drought or selection for resistance to hydraulic cavitation associated with reduced stomatal conductance and carbon assimilation in late-flowering ecotypes [31,32].

The empirical testing of the WBE model’s assumptions and predictions involves numerous challenges, primarily due to the complexity and scale of measuring branching traits in vast vascular networks. Evaluating the WBE model requires precise and comprehensive measurements of several branching traits across thousands of branches within a network, a labor-intensive and time-consuming process [33,34]. Confronting theoretical predictions with empirical data introduces uncertainties about the most effective methodologies and interpretations. While some studies have found results consistent with WBE’s predictions for an optimized branching network [35,36] others have identified discrepancies. These discrepancies are particularly notable when comparing branching traits and scaling exponents at the whole-tree level versus local branching junctions [33,37–39]. These findings suggest that variability in branch size and function within a tree crown may decouple local branching architecture from whole-tree allometries, challenging the validity of WBE’s core assumptions. Such mixed results highlight a fundamental tension between macroscopic scaling relationships and the microscopic details of vascular geometry.

Building on past work [28,34], we propose methodological advances that incorporate a more comprehensive and accurate approach to measuring and analyzing branching traits to address these challenges. We revisit two assumptions that have characterized previous tests of WBE:

First, we emphasize the importance of statistical self-similarity across the entire branching network rather than relying solely on symmetrical branching at node junctions. This approach is based on the WBE model but focuses on measures of branching traits that better capture self-similarity in branching networks. By avoiding the aggregation of point branch measures across branch nodes and levels (eg. [33,37–39]) we aim to reduce bias and improve the accuracy of scaling exponent predictions. We replicate methods based on node-level ratios to demonstrate how they can bias calculations of allometric scaling exponents [33,34,40].

Second, we address the critical assumption of terminal branch dimension ‘invariance’, which directly influences the metabolic capacity of the network [41,42]. Simplistic application of this assumption to real biological networks can lead to erroneous estimates of WBE predictions. Variability in twig and leaf size (terminal elements) at the end of branching networks is linked to variability in their physiology and metabolic rates [43]. Ignoring terminal branch variability can significantly bias attempts to measure allometric scaling relationships in trees. Instead, we account for terminal branch size variability by measuring branch cross-sectional area to estimate sapwood area as a proxy of total leaf area [44]. This approach allows us to more accurately assess WBE predictions by incorporating the physiological and metabolic variability of the terminal elements, advancing from past work assuming invariant terminal branches [33,34,40].

For finite-sized organisms, the WBE model does not predict a pure power-law but rather a curvilinear relationship between the logarithm of metabolic rate and body mass. As vascular networks grow and the number of branches increases, the network will asymptotically approach the optimal ¾ scaling relationship between total metabolic rate and total body size [27,42].

While curved metabolic scaling, called the finite size effect, has been empirically demonstrated in interspecific plant and animal studies [16,45,46] WBE’s predictions for finite size effects in metabolic scaling have not been tested within and across tree branching networks. Until recently, entire tree crowns and all of their branches have been logistically infeasible to measure due to the practical limitations of measuring thousands, if not tens of thousands, of branches

We utilize a global dataset of tree Terrestrial Laser Scanning (TLS) data and hand-measured tree branching networks to test WBE’s allometric predictions for branching traits and scaling. The extensive TLS dataset allows us to assess variability in branching traits within and across tree networks, providing a robust basis for evaluating the WBE model across diverse network sizes. We address potential biases unique to TLS data by implementing a standardized workflow that combines algorithmic and manual corrections, validated through replication with manually measured twigs and trees.

We hypothesize that allometric branching traits, accounting for heterogeneity in tree crowns, can better capture the self-similarity in branching architecture fundamental to the WBE model. This approach aligns closely with the theoretical framework and offers a more accurate test than previous methods, focusing solely on node-level traits. Additionally, we propose two key corrections to avoid biased estimates of metabolic scaling: quantifying network metabolic capacity by capturing terminal branch size variability and accounting for curvature in allometric predictions. With these corrections, we predict that diverse tree crowns will exhibit robust ¾ metabolic scaling, consistent with network branching geometry. We begin by describing how to measure metabolic scaling directly by linking the concept of metabolic capacity to total leaf area in plants. We then review WBE’s approach to plant vascular allometry and test three key predictions in TLS-derived tree models and hand-measured branching networks.

### Theory

Metabolic scaling relates the size of an individual organism to its metabolic rate. The WBE model explains how organismal metabolic rate, *B*, scales with total size, *M*, primarily through specific branching geometries of vascular networks. This model provides a theoretical basis for deriving Kleiber’s Law, which states that *B*∝*M*^*θ*^, where *θ* = 3/4. The metabolic scaling exponent, *θ*, mathematically expresses how metabolic rate changes with size.

Historically, metabolic scaling relationships have been determined empirically by regressing measures of metabolic rate against organismal mass either within a single individual across its development, within a species across individuals, or across species [15,47,48]. Metabolic capacity and metabolic rate may be distinguished as the difference between potential and realized performance in organisms [49]. Rather than more direct physiological proxies of *B* such as gas exchange, mitochondrial activity, or heat fluxes, WBE permits us to rely solely on vascular network geometry to quantify the metabolic capacity of an organism.

WBE defines the branching network’s ‘resource or metabolic capacity’ [41] as the range of resource flux through the base of the branching network that meets the variable metabolic rates of distal tissues and cells. Focusing on plant vascular networks, we can define metabolic capacity as the total leaf area *A*_*Tot*_ of a plant. Within a tree branch, total resource flux through that branch as photosynthesis, transpiration, and respiration all vary proportionally to one another across time scales, but is equal to the sum of the sap flux requirements of all distal leaves. In other words, since all plant functioning (including metabolism) is ultimately downstream of photosynthesis, the WBE model assumes that *B*∝ *A*_*Tot*_, and we refer to leaf area scaling and metabolic scaling interchangeably [27,30].

The following sections build up three previously derived allometric expressions (Equations 4a, 6, and 7), representing the WBE model’s core assumptions [27,50]. Together, they constitute the theoretical foundation for our major goal: operationalizing the assessment of WBE predictions within TLS models of tree crowns. To do this, we describe two empirical traits that can more accurately quantify metabolic scaling: i) a recently proposed proxy for leaf area (metabolic capacity) measurable solely from branching geometry [44] and ii) the relative network depth, a trait we introduce that captures the signature of curved allometric scaling due to finite size effects in branching networks. We begin by showing how the WBE network model predicts metabolic capacity as total leaf area (*A*_*Tot*_) from plant branching architecture.

### Scaling total leaf area with network branching architecture

The Pipe model of plant vascular architecture [51] and more recent elaborations are used to derive fundamental expressions linking leaf area and branch radii [27,36,52,53]. WBE defines the total leaf area supplying a network as a macroscopic measure of metabolic capacity in plants:

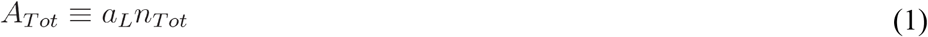

where *A*_*Tot*_ is the total leaf area of the branching network, *a*_*L*_ is the average area of a leaf, *n*_*Tot*_ and is the total number of terminal vascular tubes (tips, i.e. leaf petioles). To analyze the scaling of *A*_*Tot*_ with plant size, WBE defines a branching network with discrete hierarchical levels *N* →[0,*T*], where *N* is called the network depth. The trunk of the tree is *N* = 0, *N* = *k* is an intermediate scale, and *N*=*T* are the terminal branch tips (Box 1). is equivalent to the integer number of bifurcations (branching levels) from trunk to tip in a symmetrical network where all paths branch an equal number of times. Each branching level *k* of the network contains some number of branches *n*_*k*_ (so that *n*_*T*_ = *n*_*Tot*_ is again the number of terminal tips), and a subtree is a network fragment containing a single branch at level *k* and all distal connected branches, including *n*_*T,k*_ distal connected terminal tips.

The WBE model assumes self-similarity in the network’s geometry. As a result, the branching furcation ratio does not vary with the depth *N*. It is defined as the ratio of distal terminal tip count at successive levels of the hierarchy *n* ≡*n*_*T,k*_*/ n*_*T,k+*1_, where *n* = 2 for a bifurcating network. The furcation ratio can describe the total count of terminal tips for a subtree at *k* level as an exponential function of the network depth:*N*_*T,k*_ = *n*^*T −k*^[26,27,50]. For example, when *k* = 0,*n*_*T,0*_=2^*T*^ is the total number of tips in a tree, and k = *T,N*_*T,T*_ = 1 for a single tip. The branching ratio allows arbitrary scales to be compared within the branching network using allometric expressions. Using this ratio and Equation 1, we can calculate the distal leaf area for any branch at level *k* as:

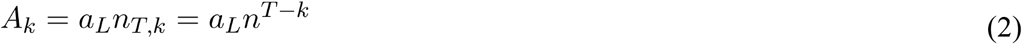

Equations 1 and 2 both emphasize that leaf area *A*_*k*_ is linked to metabolic capacity via terminal tips *n*^*T* −*k*^ at all scales in the network. The key insight of WBE comes from applying self-similarity to the physical dimensions of the branches. Specifically, the self-similar ratio between successive branch radii in the hierarchy, the radius scaling ratio, can be defined in terms of the furcation number:*r*_*k*_ /*r*_*k+*1_ ≡ *n*^*a/2*^, [27,50]. If a =1then the branching is “area preserving” and the cross-sectional area is preserved across successive branching levels throughout the network.

We can now rewrite the tip scaling term *n*^*T* −*k*^ to provide a size-based expression for distal metabolic capacity in terms of branch radii. Substituting the radius scaling ratio, we arrive at an expression for metabolic capacity in terms of *r*_*k*_:

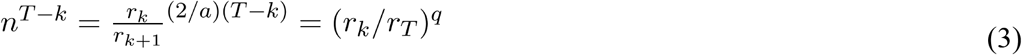

where *q* = 2/*a*. The substitution 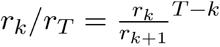 is a key step that summarizes potential variation in the node-level radius scaling ratio across a network, and expresses it allometrically with the exponent *q*, previously referred to as the conductance allometry [30,50,54]. Past work has emphasized expressing branching traits as exponents to control for heterogeneity and measure statistical self-similarity [34]. We apply it to two other allometric predictions below. This allometric formulation of node-level branching trait variation controls for heterogeneity in tree crowns while expressing possible species-level differences in branching architecture. For example, when *a* =1 the average node-level branching is area-preserving, while when *a* ≠1 the branching could be area-increasing (*a* > 1) or area-decreasing (*a* < 1) on average across the whole network. For clarity, we can restate the proportionality between leaf area, network terminal metabolic capacity, and the subtending network basal radius as follows:

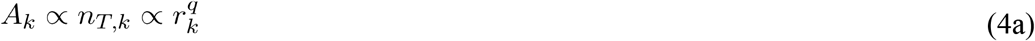

The scaling of the total leaf area of the subtree can be written in terms of an exact allometric equation for the basal subtree radius *r*_*k*_ and the distal branching geometry implied in exponent *q*:

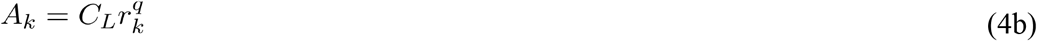

where 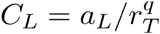, is an allometric normalization that could vary between individuals or species. The value of *C*_*L*_ denotes the direct proportionality between the average single-leaf area and terminal branch cross-sectional area [55]. This proportionality is well-known as one of Corner’s rules [56,57]. Equation 4, particularly the trait *q* = 2/*a* has been attributed to the hydraulic architecture of plant stems, especially xylem anatomy [50,51], but complementary biomechanical explanations emphasize that this is a central, multifaceted, and general relationship in plant architecture [58,59].

### Leaf area as the sum of terminal branch cross-sectional areas

Next, we show how an operationalized proxy of *A*_*k*_ allows us to measure the exponent *q* in TLS-based tree models. Network metabolic capacity in plants is determined by *A*_*k*_ across scales, which in turn covaries with terminal branch size and geometry. There is considerable variation in the size, age, and functional capacity of terminal plant organs i.e. twigs and leaves [43]. Notably, twigs are characterized by allometries that scale leaf number by total size [60–62]. Further, new methods for imaging tree networks (e.g. terrestrial laser scanning or TLS) often introduce error and variability near the terminal branches [39,63]. In practice, TLS structural models contain ‘terminal’ branches much larger than the size of twigs (usually centimeter vs. millimeter scale, respectively) and so can exhibit wider variance in size than manual measurements of branch terminals [39,63]. In short, accounting for variation in branch size is a key step toward quantifying metabolic capacity as *A*_*k*_, distal leaf area.

Equation 4a presents an important ambiguity in relating *A*_*k*_ to metabolic capacity, since ‘terminal tips’ *n*_*T,k*_ can be interpreted as leaves or twigs bearing multiple leaves, all of which can vary in size. Recent work has proposed terminal cross-sectional area as a robust metric for sapwood area estimation that better handles biases in TLS structural models [44,64]. As an improvement over counting terminal tips *n*_*T,k*_ to calculate metabolic capacity, we apply this summed cross-sectional area of distal terminal branches to account for variation in terminal branch size. Equation 4b justifies the core intuition for this leaf area proxy. If *T ′* represents an arbitrary depth for a given terminal branch in a TLS tree model, generally above the true twig scale, then 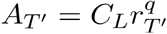, where *q* = 2 assumes that radial scaling is approximately self-similar between *T ′* and the true twig scale *T*(Box 1). Therefore, we propose an alternative expression to test as a replacement for Equation 4a, where we sum the cross-sectional area of all observed terminals distal to branch *r*_*k*_:

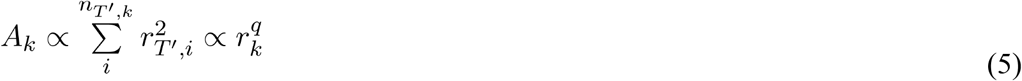

This sum of terminal cross-sectional areas serves as a clearer physiological linkage to potential variation in leaf area, and a more robust way to measure the exponent *q*. Additionally, this linkage is independent of *r*_*k*_, and accounts for variation in the geometry of terminal elements, namely *r*_*T ′,i*_. Again, we express this for arbitrary depth *T ′*because terminal branches in TLS are rarely ‘true’ terminal twigs (and never petioles) but instead represent a range of sizes and depths resulting from branch damage/senescence, instrument error, and especially truncation from data processing. This method allows *q* to vary across tree crowns, species and environments, which may reflect underlying differences in xylem architecture or other factors affecting the scaling of branch radii.

### Curved metabolic scaling

Next, we link Equation 5 with network size to reach a prediction for metabolic scaling readily applicable to TLS scans. First, we expand from network radii to include overall organismal size, using the relationship between branch radius and distal network mass *M*_*k*_

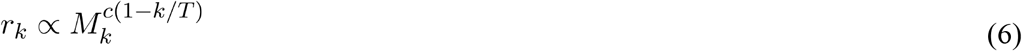

where *c* = *θ/q=*3/8 when *q* = 2, predicting covariation between these allometric exponents, discussed below. We refer to *c* as the mass-scaling exponent following previous work [30,54]. The mass-scaling exponent results from the self-similar, volume-filling nature of the branching network [27]. The mass-scaling exponent integrates the effects of branch length scaling in the mass dimension, satisfying biomechanical stability constraints [27,50,65]. By combining with Equations 5 & 6 at each branching level *k* and noting that *θ*= *cq*, a similar prediction is made for the distal leaf area, providing WBE’s core prediction for metabolic scaling in plant networks:

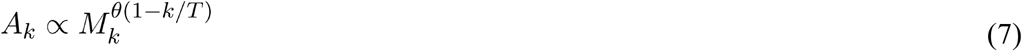

Both Equations 6 and 7 predict finite-size curvature on log-log axes. Rewritten, these equations imply that *logA*_*k*_ ∝(*θ* − *θ*(*k/T*))*logM*_*k*_ and *A*_*k*_,*M*_*k*_, *θ* covary with a relative network depth *k/T*. As subtrees increase in size (i.e. as *k* → 0), then *logA*_0_ ∝(*θ* − *θ*(0*/T*))*logM*_0_ so that *logA*_0_ ∝ *θlog M*_0_ for the whole tree, given *T* is large [42]. Assuming constant tissue density, the total network volume scales similarly, predicting 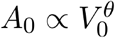. This provides a method for evaluating finite-size curvilinear metabolic scaling within and across large branching networks, such as mature trees.

An underexplored consequence of this theory is that variation in metabolic scaling can be described by the exponents in Equations 5 and 6, such that *θ = cq* when networks are large and deeply bifurcated. Describing variation in metabolic scaling as the product of these two quantities, termed conductance or water-use scaling *q* and mass scaling *c*, has been emphasized previously [30,50,54] but not evaluated thoroughly within entire tree crowns for diverse species and sizes. By directly evaluating the allometric branching traits *q* and *c*, we quantify metabolic scaling without aggregating node-based, symmetrical branch scaling ratios [33,37–39]. This approach captures covariation between allometric branching traits *q* and *c* arising from architectural differences in tree crowns.

In this section we described two empirical traits for measuring metabolic scaling in tree branching networks. Equation 5 is a prediction for metabolic capacity scaling relying solely on branching geometry as a proxy for leaf area. We introduce a novel operationalization of the quantity *k/T* which we refer to as relative network depth (see Methods), allowing us to control for curvature within and across tree crowns and more accurately measure *c* and *θ*.

Next, we test the three key allometries in Equations 5, 6, and 7, which include the allometric branching traits *q* and *c* expressing core biophysical assumptions of WBE. By comparing scaling in trees processed algorithmically and manually, we aim to reduce bias in branch geometry at small sizes. Additionally validating against manually measured plant networks helps distinguish systematic bias in TLS models from natural variability in tree architecture scaling. We demonstrate that these processing steps are essential for revealing convergence in allometric exponents toward WBE predictions in TLS data.

## Methods

### TLS tree models

We compiled TLS data for 339 trees, mostly from the published literature (Table S1). These data comprised point clouds of individual trees split into two groups. The first group included 62 trees from Cameroon [38,66] and 32 trees from French Guiana (author data). We analyzed these two datasets with a standardized cleaning methodology that controls for bias in fine branching geometry. The second group of trees represented a variety of methodologies from many studies without additional processing steps such as manual editing, statistical downsampling, or de-noising. However, some interventions were performed in their respective published sources. Many trees were leaf-off or defoliated, but other foliated trees were processed leaf-on.

From the point clouds, we constructed Quantitative Structure Models (QSMs) for each tree using the TreeQSM (v2.4.1) software package [67] in MatLab 2019b. QSMs provide volumetric models of tree point clouds by fitting a network of connected cylinders to branching geometry. We performed a series of QSM fits using the ‘treeqsm’ driver function. Parameter ranges were set based on previous recommendations [63,68], and optimal QSMs were extracted with the ‘select_optimum’ function. TreeQSM optimization criteria were selected for each set of tree clouds independently by visual inspection to account for differences in scanning conditions across datasets (Table S2). Visual inspection threw three exceptionally poor tree models out of the analysis. Ultimately we successfully fit 246 cylinder models from over 40 tree species in tropical and temperate sites in the broader group from the literature (see Table S2 for a list of individuals with metadata).

Internodes in the QSMs are segments separated by any branching point in the hierarchy of connected cylinders. TreeQSM models each internode as a sequence of cylinders to accommodate branch taper and shifts in branch orientation. To calculate the length of an internode, we summed the length of individual cylinders within each segment. Since the radius of these cylinders also varies, we took the average of the radii across all cylinders within an internode. We made these calculations and further analyzed the QSMs, in the R programming language [69].

### TLS and QSM cleaning protocol

To validate predictions for allometric branch scaling, we tested for the impact of bias in different processing workflows on observed scaling relationships. Cleaning is the addition of manual, visual correction steps to algorithmic outputs of TLS tree processing (Figure S6). This semi-automated workflow reduces the impact of biases known to affect QSMs [63]. To illustrate the actual effect of cleaning models, we compared cleaned versions of trees to biased structural models of the same individual tree scans without cleaning steps. We also compared these results to the broader, biased global database of TLS scans.

The cleaning process specifically targets biases resulting from inflation of branch radius and length at small sizes (near one centimeter) and overfitting branches to noisy point cloud data. This process is labor-intensive but combines software-based algorithmic and manual processing steps for: i) separation of leaf and stem tissue and ii) fitting cylindrical branching geometry to point clouds. Defoliation was accomplished using LeWos [70]. Simply applying defoliation algorithms in an automated way does not generally improve analyses of branching architecture due to misclassification (both Type I & II errors), which exacerbates errors at small branch sizes.

Manual correction of defoliation errors in CloudCompare is the key step for effective tree model cleaning in the data presented. After QSM fitting in TreeQSM, cylinders are manually corrected for branch diameter, orientation, and connection errors. This was accomplished in AMAPstudio-Scan [71].

### Ground-truthing with hand-measured plant network data

In addition to cleaning, we performed validation by testing the same predictions for branch allometries in a collection of manually measured tree networks. The diverse networks cover eleven angiosperm and gymnosperm species from tropical and temperate areas (Table S4). The data also span a significant size spectrum, from millimeter-scale branches to a dataset of large, felled tropical trees with branches manually measured from trunk to 10cm diameter. This allows the assessment of scaling patterns across a range well below TLS resolution (< ∼1cm) to individuals rivaling the largest TLS scans in our analysis. We measured the same key allometric scaling relationships at the subtree scale for each manually measured tree network with SMA regressions.

### Subtree scaling regressions for measuring branch allometry

We assessed the allometric predictions of the network model within tree crowns through regressions on subtrees. Subtrees were defined by a basal internode and included all distally connected internodes. This allowed us to identify a set of terminal internodes and measure each subtree’s total volume (*V*_*k*_). We measured the subtree volume as the sum of distally connected cylinder volumes defined by the QSM. We measured metabolic capacity in a subtree either as: i) the integer number of terminal internodes, or ii) the total cross-sectional area of terminal internodes.

Bivariate regressions were conducted by fitting standardized major axis regressions to log-transformed architectural data with the ‘smatr’ package in R [72]. Major axis regressions model error in the predictor and response variables, the true values of which are unknown in our study [73].

We also estimated leaf area scaling with a curved model using nonlinear least-squares regression on the model *logA*_*k*_ = *logV*_*k*_ * *a* *(1−*b*\*(*N*_*k*_))+*c*. This model directly implements Equation 7, where the parameter estimates the leaf area scaling exponent. Most importantly, it incorporates the relative subtree depth 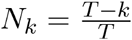 to assess and control for a key WBE prediction: curvature within and across branching networks networks. We assess curvature by analyzing the magnitude and significance of the coefficient *b*.

The relative network depth should range *N*_*k*_→[0,1] from trunk to tip. We reverse the indexing from the network theory presented above and set the depth of the terminals so that *k =* 0, and the depth of the whole tree network from the trunk is considered *k* = *T*. To estimate *k* for an arbitrary subtree, we reverse indexing so that *n*^*T*−*k*^*=N*_*T,k*_ instead becomes *n*^*k*^=*N*_*T,k*_. Subtree network depth is then estimated as *k* = *log*(*n*_*T,k*_)/*log*(2), where we assume the branching ratio. This effectively assumes that each subtree path within a crown bifurcates the same number of times, so tips and bifurcations are well-correlated [46]. The above logarithmic expression tends to be closely correlated with the maximum order index produced by other branch ordering schemes, and previous work has pointed out that branch ordering may be highly misleading to calculate depth *k* in asymmetrical networks [5,74–76]. When fitting the curved expression interspecifically across all subtrees (Figure 4), we set the maximum depth *T*_*max*_ to the maximum depth across all trees (i.e. the depth of the largest tree) to get the best estimate of the asymptotic limit at the interspecific scale. We also tested Equation 6 with this method.

To better illustrate curvature, we also fit the model *θ*_*LA*_ = *C*−*Re*^*Kn*^ across individual tree crowns, where *θ*_*LA*_ is the straight-line (SMA) leaf area scaling exponent, *N* is the maximum depth for a given crown (not the interspecific *T*_*max*_), and *C,R,k* are fitted constants. We used an identical model to analyze variation in the mass scaling exponent across individual tree crowns.

We used the base R function ‘nls’ to fit curved allometries to log-transformed data. When constraining the curvature parameter to fit mass allometries, we selected the algorithm ‘port’; otherwise, we used the default setting, a Gauss-Newton algorithm. 95% confidence intervals were calculated with the ‘confint2’ function in package ‘nlstools’ [77].

### Closed form theorems from scaling ratios

We replicated node-level methodologies, and computed theoretical metabolic scaling exponents to compare with our approach. Branch scaling ratios were computed at each branching node using the dimensions of fitted cylinders. The radius scaling ratio, *β* ≡ *r*_*k+*1_/*r*_*k*_, is defined as where *r*_*k*_ is the radius of a branch at level in the network (reciprocal of the ratio of radii in Equation 3). The length scaling ratio, *γ*, is defined as *γ* ≡*l*_*k+*1_/*l*_*k*_ where is the length of a branch at level *k*. At each bifurcation *β* and *γ* are computed for each connected branch. WBE predicts that for a symmetrical space-filling and area-preserving network *β* and *γ* will equal *n*^−1/2^ and *n*^−1/3^ respectively. The metabolic scaling exponent is derived as:

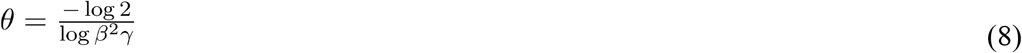

This expression for the metabolic scaling exponent was calculated for each tree using average values of *β* and *γ* calculated across all connected branches within a given tree. Whole-tree averages were calculated from nodes with scaling ratios less than 1. Across plant morphological datasets, child branches with dimensions greater than parent branches frequently occur. In our structural models, approximately 25% of branching nodes have child branches that exceed parent dimensions in either length or radius. This can lead to undefined behavior in Equation 8 because when *β*^2^*γ* > 1, the exponent becomes negative–this behavior has led to filtering nodes in previous publications [34]. Subsequent work also showed that alternative equations for metabolic scaling not requiring data filtering did not show qualitative differences in branching traits or metabolic scaling exponents [46], so we replicate the simplest method to illustrate the flaws in the node-level approach in general.

### Data analysis

Our analyses emphasize two comparisons to test WBE predictions for allometric scaling. First, we analyze scaling within individual tree crowns and also fit a single regression model to all network geometry across all sampled individuals to infer an interspecific relationship.

Second, we fit separate models for biased and unbiased data, depending on the processing steps applied to individual tree scans. These two groupings are summarized in Table 1.

**Table 1:**
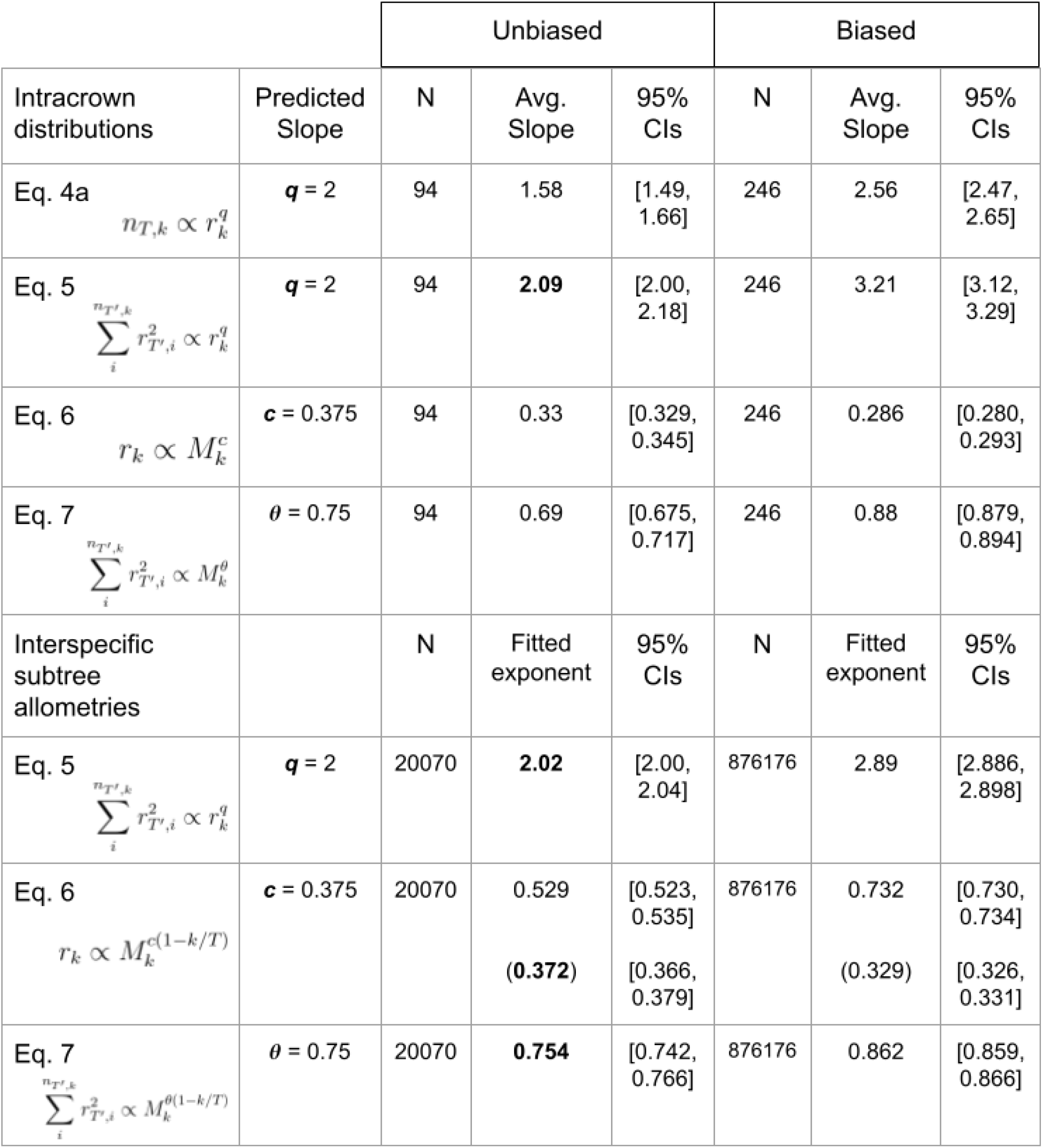
Summary of regression experiments in two groups: (i) Models fit with straight lines within individual tree crowns (modified Equations 6 and 7 to ignore curvature), and (ii) models fit for all subtrees across all individuals (‘Interspecific subtree allometries’), using curved models where applicable. Exponent measurements in bold significantly overlap theoretical predictions with one-sample t-tests for intracrown distributions, or fitted parameter values in SMA/NLS regressions for interspecific subtree allometries. For interspecific results on Equation 6, slopes in parentheses are alternative fits with constrained curvature.

|  |  | Unbiased |  |  | Biased |  |  |
| --- | --- | --- | --- | --- | --- | --- | --- |
|  |  | N | Avg. Slope | 95% CIs | N | Avg. Slope | 95% CIs |
| Intracrown distributions | Predicted Slope |  |  |  |  |  |  |
| Eq. 4a<br>$n_{T,k} \propto r_k^q$ | **q = 2** | 94 | 1.58 | [1.49, 1.66] | 246 | 2.56 | [2.47, 2.65] |
| Eq. 5<br>$\sum_i^{n_{T',k}} r_{T',i}^2 \propto r_k^q$ | **q = 2** | 94 | **2.09** | [2.00, 2.18] | 246 | 3.21 | [3.12, 3.29] |
| Eq. 6<br>$r_k \propto M_k^c$ | **c = 0.375** | 94 | 0.33 | [0.329, 0.345] | 246 | 0.286 | [0.280, 0.293] |
| Eq. 7<br>$\sum_i^{n_{T',k}} r_{T',i}^2 \propto M_k^\theta$ | **θ = 0.75** | 94 | 0.69 | [0.675, 0.717] | 246 | 0.88 | [0.879, 0.894] |
| Interspecific subtree allometries |  | N | Fitted exponent | 95% CIs | N | Fitted exponent | 95% CIs |
| Eq. 5<br>$\sum_i^{n_{T',k}} r_{T',i}^2 \propto r_k^q$ | **q = 2** | 20070 | **2.02** | [2.00, 2.04] | 876176 | 2.89 | [2.886, 2.898] |
| Eq. 6<br>$r_k \propto M_k^{c(1-k/T)}$ | **c = 0.375** | 20070 | 0.529<br>**(0.372)** | [0.523, 0.535]<br>[0.366, 0.379] | 876176 | 0.732<br>**(0.329)** | [0.730, 0.734]<br>[0.326, 0.331] |
| Eq. 7<br>$\sum_i^{n_{T',k}} r_{T',i}^2 \propto M_k^{\theta(1-k/T)}$ | **θ = 0.75** | 20070 | **0.754** | [0.742, 0.766] | 876176 | 0.862 | [0.859, 0.866] |

Within individual tree crowns, we used paired t-tests to evaluate differences in conductance allometry results for two measures of metabolic capacity (Equations 4a and 5) and compared these between biased and unbiased data.

We also tested WBE’s theoretical predictions for distributions of scaling exponents calculated within individual tree crowns using one-sample t-tests separately in biased and unbiased data. We tested each focal allometry in both groups (Equations 4a, 5, 6, and 7). We compared the variability of the conductance and mass-scaling distributions to the WBE prediction for covariation in those quantities (*θ*=*cq*).

At the interspecific scale, we pooled all individual subtree measurements and extracted single regression models separately in biased and unbiased data for the focal allometries related to metabolic scaling (Equations 5, 6, and 7).

Finally, we compared theoretical metabolic scaling exponents calculated from node-level branching traits to regression-based exponents for biased and unbiased trees. We fit a simple linear model to both sets of trees to evaluate how well theory-based metabolic scaling exponents correspond to the regression-based formulation of metabolic scaling using allometric branching traits.

## Results

We find that the two measures of metabolic capacity yield different predictions for the conductance scaling relationship (Equation 4a vs Equation 5). Conductance scaling exponents are significantly different in both biased (p < 0.01) and unbiased (p < 0.01) tree crowns, with the cross-sectional area proxy yielding steeper exponents than tip counting (Figure 1, Table 1).

**Figure 1:**
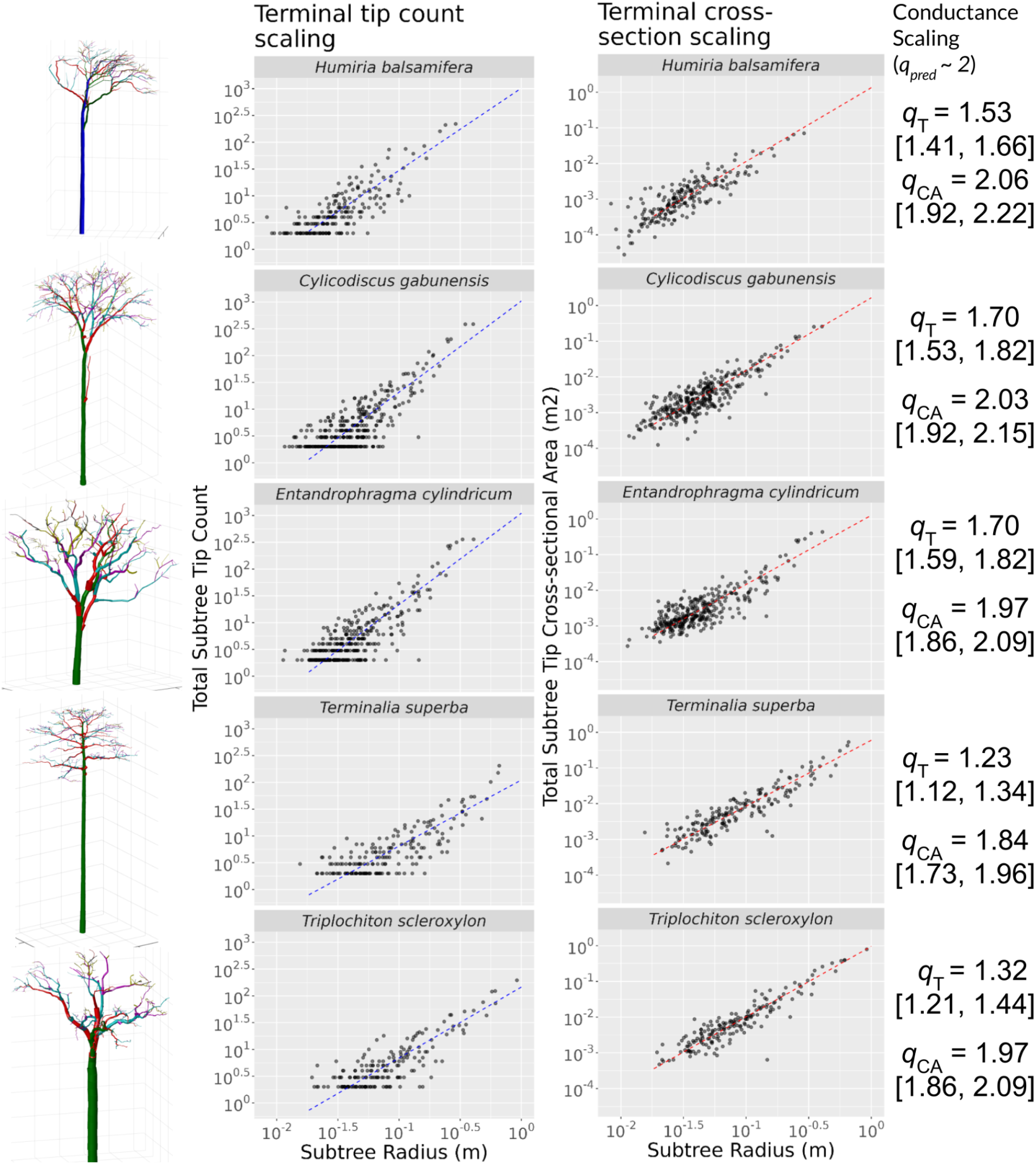
Total terminal cross-sectional area is a more accurate estimate of metabolic capacity than counting terminal branches, which yield shallow regressions due to wide variation in basal radius at terminal ends of the network. *Left)* Cylinder models derived from TLS scans of five tropical tree individuals. *Center)* Regressions on two estimates of subtree distal leaf area: Integer tip count versus terminal cross-sectional area. Dashed lines are bivariate standardized major-axis fits to Eqs. 4a & 5. Note that cross-sectional area re-scales the vertical axis by varying the terminal dimension with metric size. *Right)* Fitted values of the conductance allometry for terminal tips (q_T_) and terminal cross-sectional area (q_CA_) regressions. The two measures of metabolic capacity are significantly different in all individuals.

Average scaling exponents for models calculated within individual tree crowns vary broadly (Figure 2, Table 1). In unbiased trees, exponents for individual tree crowns significantly overlap with the theoretical prediction for conductance scaling while being significantly lower for the mass and leaf area allometries. In the broader global dataset of biased trees, the distribution of scaling exponents for individual tree crowns is significantly different from the WBE optimal prediction in all three allometries.

**Figure 2:**
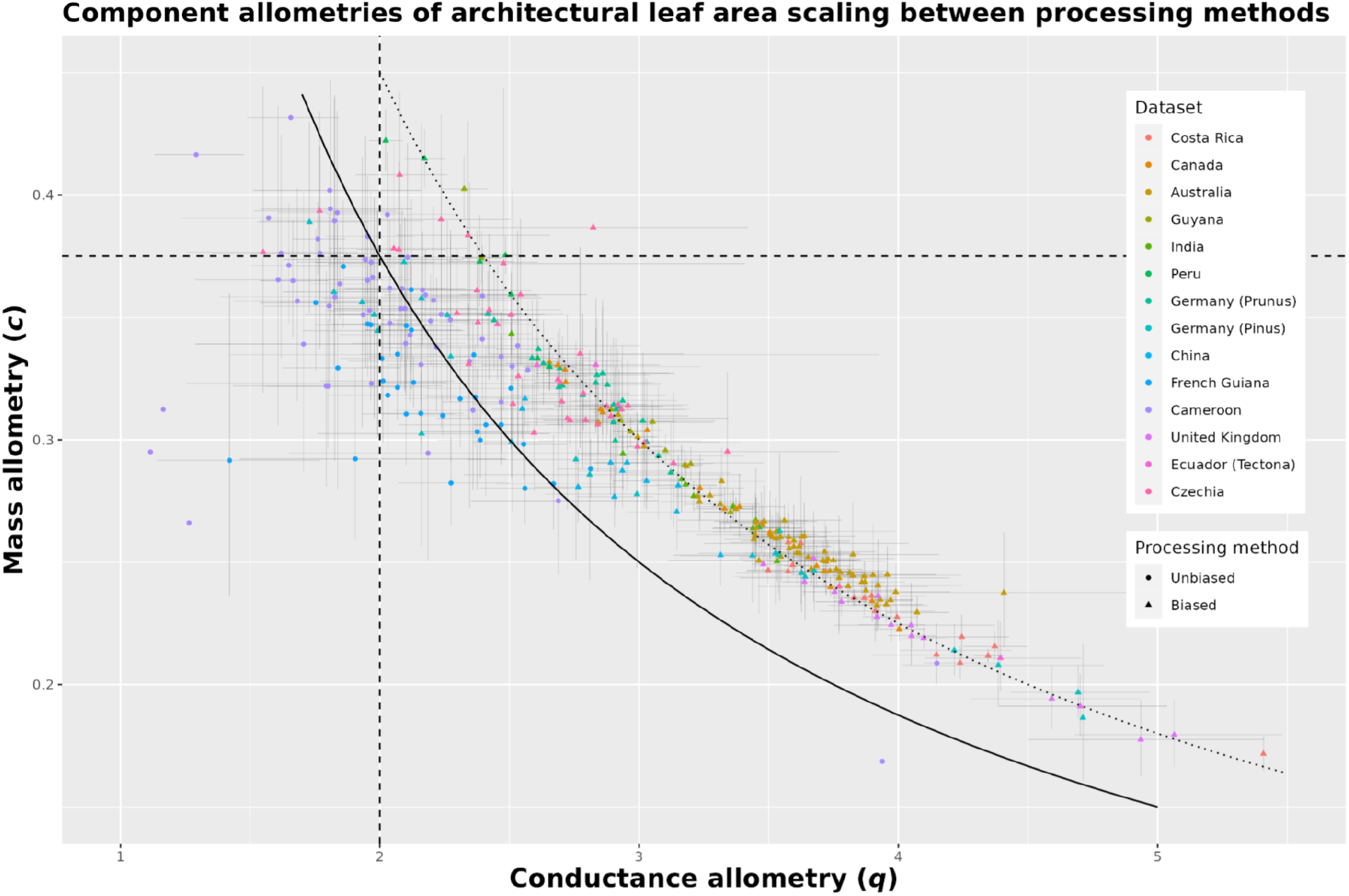
WBE predictions are supported by unbiased trees for two allometric components of leaf area scaling. Conductance scaling exponents (*q*) corresponding to Equation (5) are on the x-axis, and mass scaling exponents (*c*) corresponding to Equation (6) are on the y-axis. 95% confidence intervals. Dashed straight lines show the theoretical predictions for each allometric exponent. The curved lines are the theoretical predictions for architectural metabolic scaling exponents when values of (*q, c*) are multiplied together according to theory underlying Equation (7), where *θ* = *cq*. In the solid line *θ* = 0.75, and in the dotted line *θ* = 0.9

Unbiased trees displayed significantly different conductance scaling exponents from the same individuals analyzed prior to the cleaning process (Figure 3). The uncleaned trees were more comparable to the global distribution of conductance scaling exponents for tree models drawn from the literature.

**Figure 3:**
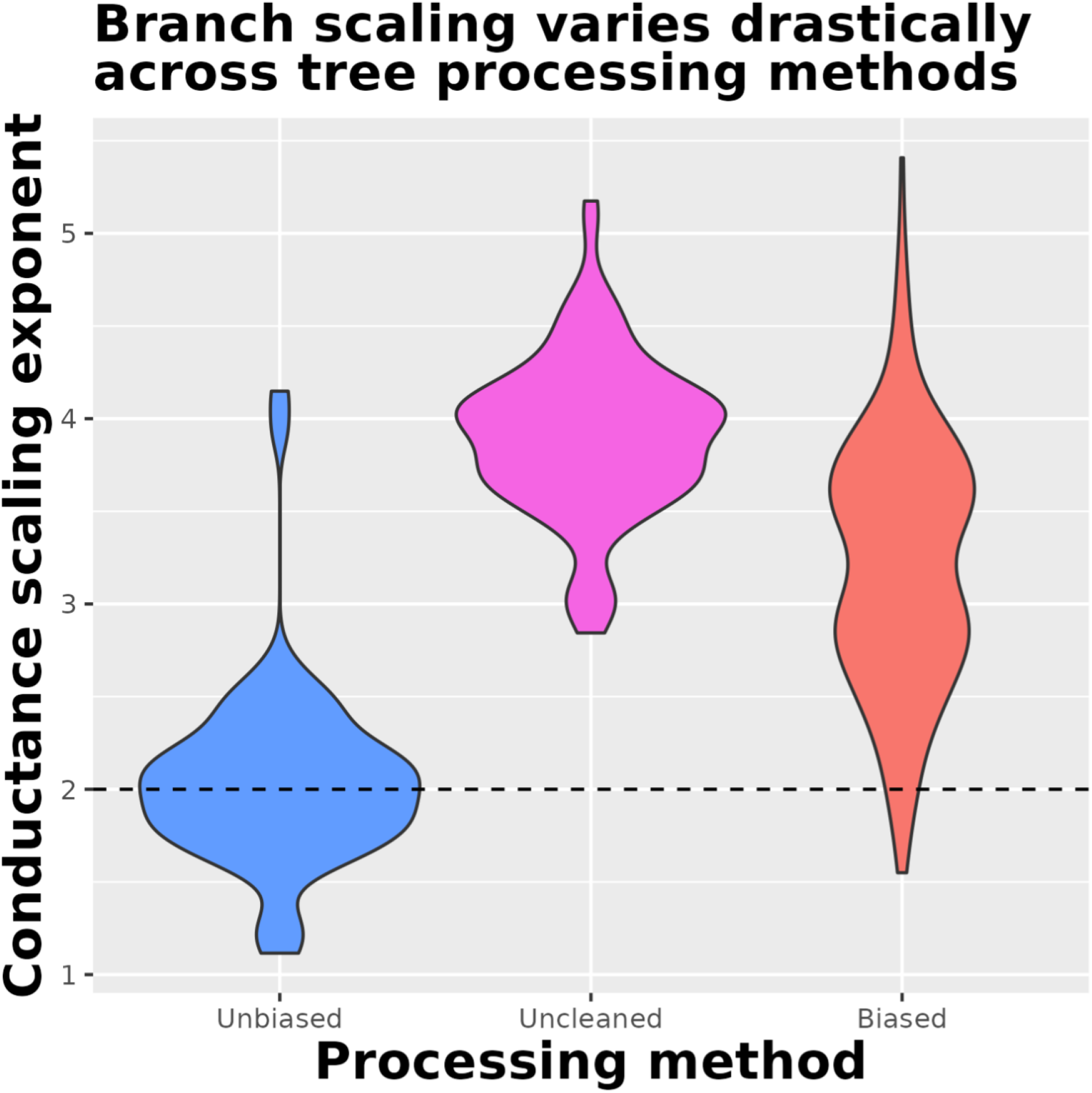
Biased tree models drastically overfit cylinders, especially at smaller size classes, compared to unbiased trees which better reflect allometric predictions. Bias is determined by the QSM processing method. Violin distributions compare conductance scaling exponents across datasets (cleaned in blue vs. uncleaned in magenta and red). Blue distribution shows conductance scaling from unbiased clean versions of the 94 tropical trees in our focal dataset. The same 94 trees were analyzed before correcting raw data for bias, and the magenta distribution shows their drastically higher distributions of conductance scaling exponents. The degree of bias matches that of the broader global distribution of tree models shown in red.

**Figure 4:**
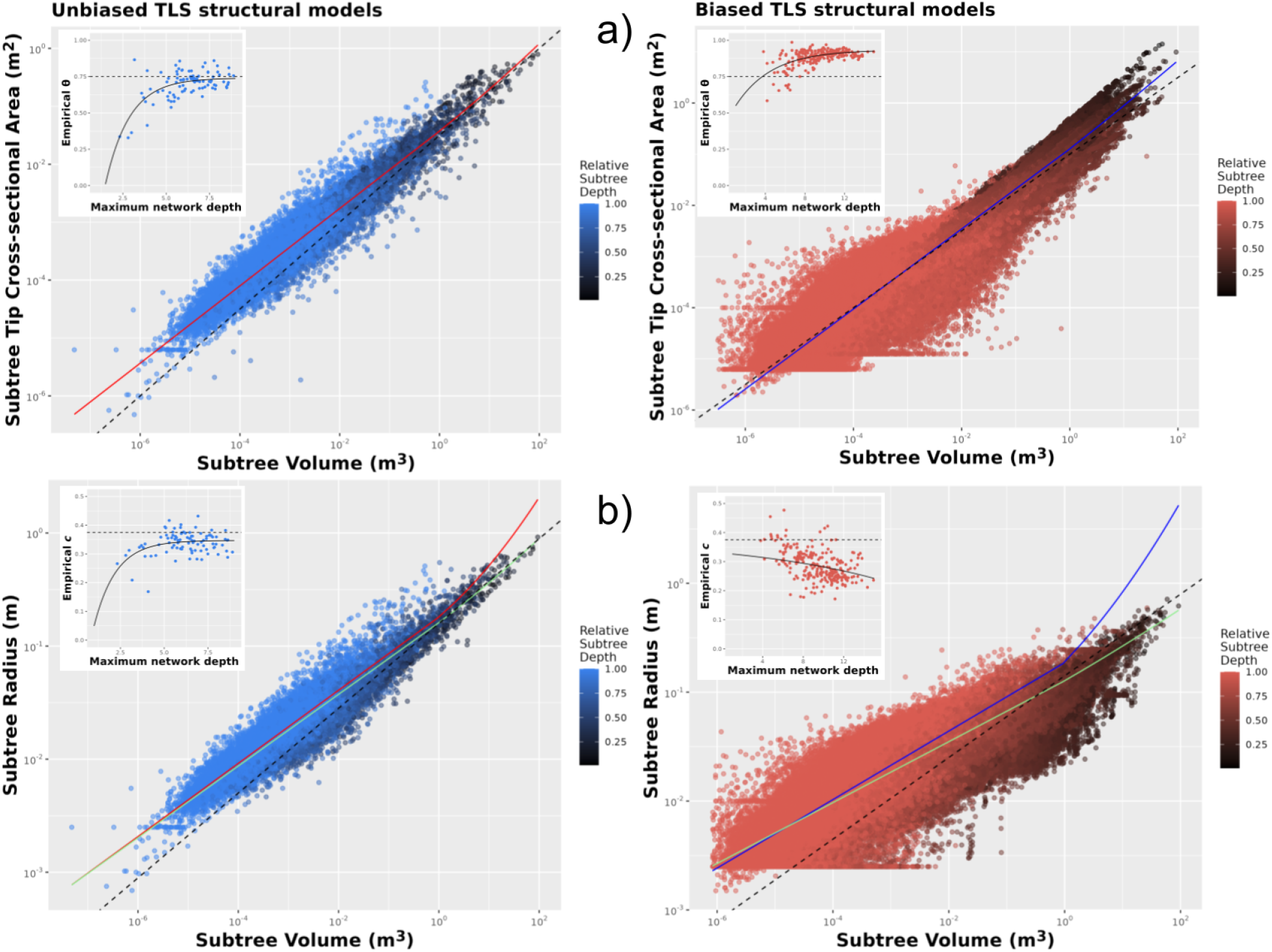
Interspecific allometric scaling shows distinct curvature in both datasets analyzed. Main scatterplots include all subtrees for each individual to maximize size spectrum and illustrate convergent patterns. **a)** Unbiased structural models support a curved model that asymptotes to the predicted metabolic scaling exponent (***θ*** = 0.754 [0.742, 0.766]). Biased structural models show distinct variation in subtree radius at smaller size classes, leading to steeper estimates of metabolic scaling (***θ*** = 0.862 [0.859, 0.866]). Dashed lines are the asymptotic ¾ prediction for leaf area scaling. **b)** Mass scaling regressions display best-fit lines with unrealistic curvature in red, so alternative fits with constrained curvature are plotted in green. Unbiased data asymptotes to the predicted mass scaling exponent (***c*** = 0.372 [0.366, 0.379]), while biased data deviates below it (***c*** = 0.329 [0.326, 0.331]). Dashed lines are the asymptotic ⅜ prediction for mass scaling. **Insets:** Another view of curvature at the scale of individual tree models. Points are individual tree crowns, with maximum network depth and straight-line SMA fits to Equations (6, 7). Here, curvature arises from underestimating scaling relationships when fitting straight-line models to curved allometries within smaller individual tree crowns, leading to correlation with maximum network depth.

Interspecific subtree scaling analyses are consistent with all three major WBE predictions associated with metabolic scaling (Equations 5, 6, and 7). Conductance scaling across all subtrees was not significantly different from the theoretical prediction, the only allometry that agrees with the intracrown distribution (Table 1).

We report two interspecific estimates of leaf area scaling: the curved allometric coefficient and the asymptote *C* serve as interspecific estimates of the leaf area scaling exponent. Each asymptotically converges on the ¾ prediction in unbiased trees while overestimating metabolic scaling in biased tree models (Figure 4a).

The curved leaf area scaling exponent *a* = 0.862 for biased trees, [0.859, 0.866]; for unbiased trees,, [0.742, 0.766] (Table 1). Curvature *b* is statistically significant (p < 0.001) in both cases. The asymptote C is 0.933, [0.900, 0.967] for biased trees, and 0.74, [0.704, 0.775] for unbiased trees. In both cases, the asymptote *C* is comparable to the interspecific curved exponent obtained by fitting all subtrees together.

Biased trees tend toward a higher maximum network depth, likely due to overfitting cylinders, which adds extraneous bifurcations and erroneous terminal cylinders. As the average number of bifurcations increases with the total cylinder number, the scaling exponents tend to asymptote (Figure 4, insets). After ∼5 branching levels or average bifurcations along a branch path, the exponents tend toward a maximum value.

We replicated this approach for the mass scaling exponent to evaluate the effect of curvature in this allometry (Equation 6). The mass scaling exponent was not significantly different from the theoretical prediction at the interspecific scale in unbiased trees, but only when curvature is constrained in the nls fit (Table 1, Figure 4b). In this alternative fit we constrain the curvature so that *b* = 0.15, resulting in a fit where *c* = 0.3729, [0.366, 0.379]. This provides a better visual fit to the data and a comparable residual sum of squares (RSS = 1541, compared with RSS = 1339 for the best-fit line), and supports the WBE prediction for subtree mass scaling. For the fit that minimizes RSS, the mass-scaling exponent *c* = 0.5293, [0.523, 0.525] and curvature is significant. The curved mass scaling exponent *c* = 0.7328 in biased trees, [0.731, 0.734], with curvature. When curvature is constrained as above so that *b* = 0.15,*c =*0.3291, [0.326, 0.331] and provides a better visual fit as in unbiased trees.

When plotting straight-line exponents fit across individual tree crowns, the asymptote is below the expectation (*C* = 0.347, [0.332, 0.361]) but agrees relatively with the fit using constrained curvature. Fitting the curved model to straight-line exponents, the mass scaling exponent does not asymptote as expected and is negatively related to bifurcation depth.

We found a correlation between metabolic scaling exponents calculated from nodel-level branching traits, and exponents measured by allometric branching traits within individual tree crowns (Figure 5). Only unbiased trees show a significant positive correlation (R^2^ = 0.19,*β*_1_=0.495, p < 0.001) with theory-based predictions for the metabolic scaling exponent.

**Figure 5:**
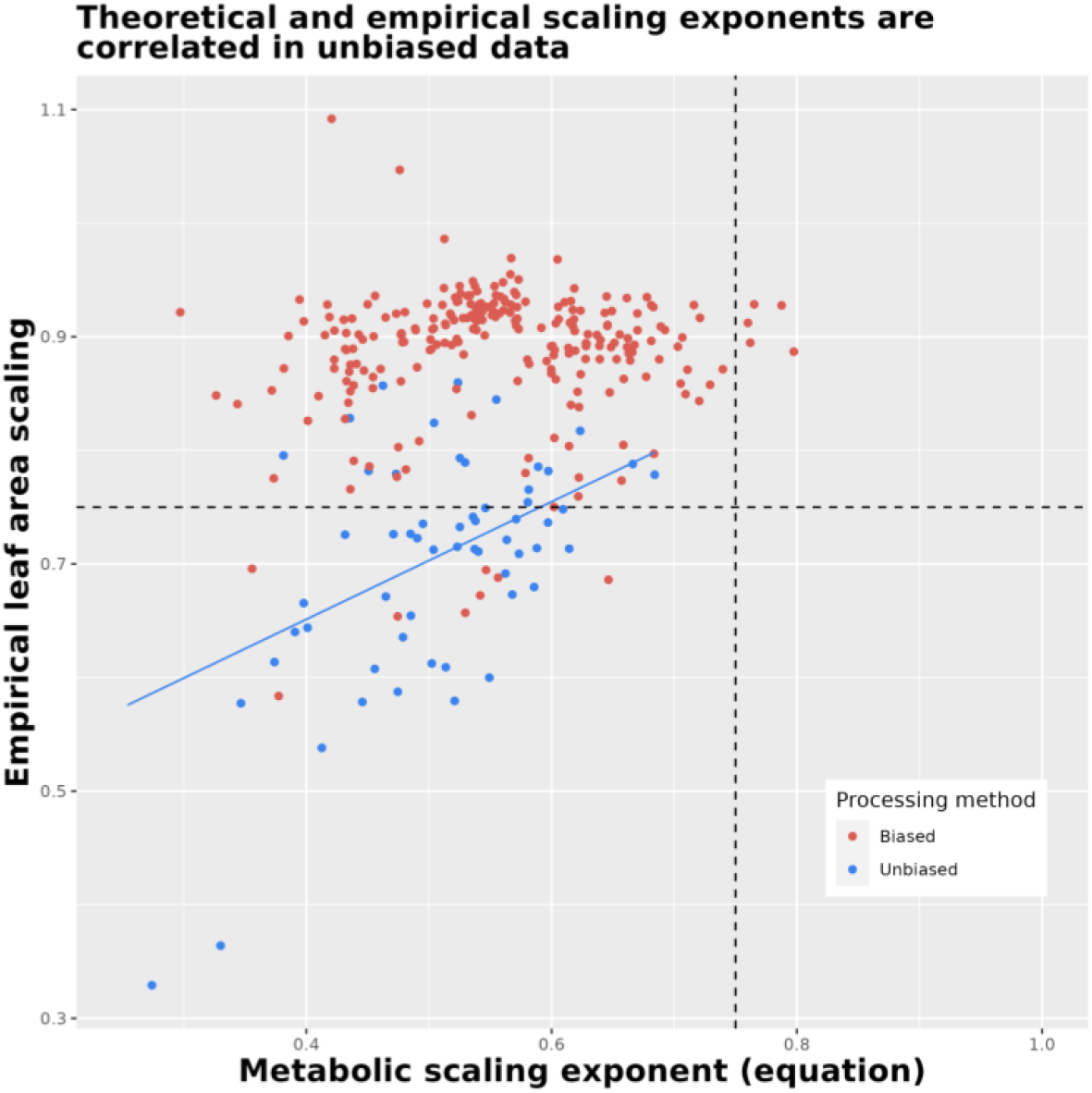
Empirical leaf area scaling plotted against the theoretical prediction for metabolic scaling calculated from theoretical, node-level branching ratios. Points are values for individual tree crowns across the two datasets. Only unbiased tree models exhibit a significant correlation. Dashed lines are the predicted optimal ¾ exponent for metabolic scaling.

## Discussion

The West, Brown, and Enquist model sheds light on tree architectural diversity by providing a mechanistic explanation of metabolic scaling and numerous other allometric relationships. WBE predictions lead to an integrative understanding of plant form by linking size and function across species through branching traits of the vascular network. We aimed to assess the assumptions of WBE using key allometric scaling predictions that describe potential variation in metabolic scaling and underlying branching traits. This study advances understanding of metabolic scaling by demonstrating how to control for various forms of bias in plant network data, and more robustly operationalize WBE theory. By utilizing Terrestrial Laser Scanning (TLS), we can now more rapidly measure metabolic scaling exponents and other key allometric traits with higher confidence in real trees.

We focused on three key components to improve on previous efforts to test the WBE model: (i) quantifying variation in network metabolic capacity by linking terminal branch cross-sectional area with leaf area variability; (ii) better capturing self-similarity by formulating allometric branching traits applicable across whole networks, rather than relying on node-level traits; and (iii) accounting for finite size effects (curvature in key allometric predictions). This approach has produced the first analysis that, relying solely on plant branching geometry, has revealed curved metabolic scaling and interspecific convergence to the canonical ¾ prediction. Our results are consistent with WBE’s core hypothesis: selective pressure on the relative scaling of metabolic capacity with network size causes specific proportionalities in vascular network geometry.

Based on plant hydraulic theory, we applied a proxy of metabolic capacity, the terminal cross-sectional area of branches. We predicted that it is proportional to distal leaf area in plant networks [44]. Metabolic capacity is linked to the geometry of terminal elements, but the specific metric must reflect WBE’s optimality assumptions and the average conditions an organism experiences. As suggested by Equations 2 & 3, counting terminal tips has been a common measure of metabolic capacity [33,34,40] but our results suggest that this method yields consistently biased underestimates of distal leaf area. Counting terminal tips often results in significant variability in distal leaf area for a given basal radius, leading to a shallow regression line and biased allometric predictions (Figure 1). This variability arises from terminal branch size variation, which generally corresponds to twigs and larger branches, rather than petioles as WBE assumes (Box 1). Consequently, in manually measured plant network fragments and coarse TLS models, the number of terminal branches is not directly proportional to the distal leaf area, producing conductance scaling exponents inconsistent with previous findings [36,54]. This result generalizes to manually measured plant network fragments, indicating it is not strictly an artifact of TLS data (Figure S1a). Instead, the total terminal cross-sectional area resolves this allometric bias by accounting for variation in size near the terminal ends of a network to better quantify leaf area and metabolic capacity. This approach improves our ability to test WBE predictions and enhances our understanding of the relationship between plant architecture and metabolic scaling.

We hypothesized that allometric branching traits robustly quantify self-similarity in tree crowns, leading to better tests of WBE’s allometric predictions. Two underlying allometric branching traits decompose metabolic scaling into distinct axes of variation: conductance scaling and mass scaling, corresponding to Equations 5 and 6. We found that individual tree crowns display notable variability in these allometric branching traits comparable to recent analyses of these allometries in tree crowns from TLS data [64]. Intriguingly, WBE predicts this pattern of variation–our data suggest these two exponents covary along manifolds defined by the metabolic scaling exponent in the manner predicted by WBE (Figure 2). This pattern represents a dimensional constraint in tree architecture that narrows the range of leaf area scaling observed across trees and may describe a trade-off between hydraulic strategies encapsulated in the conductance exponent and structural or biomechanical variation encapsulated in the mass scaling exponent. However, branch allometries are systematically biased when TLS models are not manually validated, preventing interpretation of this variability within the broader dataset of biased tree scans collected from the literature.

We focus on interpreting results from the unbiased dataset of QSMs, which align with the scaling of manually measured plant networks Figure S1b). Cleaning and manually validating structural models derived from TLS is essential for studying branch allometry given current algorithmic workflows. Experiments with digital pruning help illustrate how bias in small branch sizes is related to TLS processing workflows, and we recommend relying exclusively on cleaned trees without digital pruning in studies of branch allometry using TLS (Figure S2). Point cloud quality is a primary determinant of QSM quality in TLS studies [78]. Uncertainty from TLS data and the QSM fitting process present obstacles to modeling the smallest branches in tree crowns [39,63]. We showed that biased structural models lead to erroneous allometric estimates, particularly in conductance allometries (Figure 3, see also QSM supplement). These results on QSM biases illustrate how scaling theory validates QSM quality and reveals the holistic effects of different TLS processing workflows [44,64].

Unbiased tree crowns showed conductance scaling exponents clustered around the theoretical expectation for area-preserving branching where *q* ≈ 2. This result is evidence that total cross-sectional area of terminal branches provides a useful measure for distal leaf area in agreement with pipe-model theories of plant form [52]. These conductance exponents are consistent with average area-preservation despite node-level variability in radius scaling across tree networks. Similarly, the mass scaling exponent *c* summarizes heterogeneity in node-level branching traits across a network. When measured within unbiased individual tree crowns, these scaling exponents deviate from the ⅜ predicted by theory. This deviation from the mass scaling exponent has also been observed in studies of biomass scaling in trees and forests, although such studies rely on gross height-diameter allometries rather than branch-level analyses of entire tree crowns [79,80]. Deviation in mass scaling could cause the corresponding deviation below ¾ in the individual crown metabolic/leaf area scaling exponents. However, both these relationships are subject to curvature and are underestimated by straight-line regressions.

We evaluated WBE’s prediction for curvilinearity in the relevant scaling relationships (Equations 6 & 7). We hypothesized that curvilinearity modulates metabolic scaling in large branching networks and predicted that accounting for curvature is a prerequisite to detecting the 3/4 prediction for metabolic scaling. Interspecific analyses support the presence of curvature, and accounting for curvature is necessary to receive significant support for the ¾ scaling prediction. Curvature can be difficult to detect and constrain within individual tree crowns compared to interspecific analyses, and it can produce exponents that are not qualitatively different from straight-line fits (Figure S3). However even when accounting for curvature, the marked variation in these allometries across individuals within both manually validated network fragments and unbiased TLS models points to considerations for allometric diversity beyond the current WBE framework.

Curved allometries arising from finite-size effects may produce variation across individual tree crowns because many individuals have not developed fully toward asymptotic size limits, and these limits also vary based on functional dimensions of life history across species [81–83]. Likewise, the mass scaling exponent is a biomechanical limit that constrains metabolic scaling; in large branching networks we may expect to see exponents approaching but rarely exceeding their predicted values. To understand how ontogeny and asymptotic height growth are related in trees, the amount of forking in a tree crown has been proposed as a morphological signal of development that may influence scaling predictions and could serve as a more principled ontogenetic measure of network depth [84,85]. Future efforts measuring metabolic scaling in morphological data should prioritize accurately quantifying maximum network depth and bifurcation rates both as species-level traits and as a control on the developmental status of a given tree [38]. Still, if curved allometries arise from node-level scaling ratios varying systematically within branching networks rather than remaining constant as WBE assumes, self-similarity assumptions in branching architecture may need to be more carefully qualified and applied.

### The status of self-similarity in WBE

The allometric branching traits we rely on here, formulated as scaling exponents, help overcome the problem of heterogeneity within tree crown geometry and capture statistical self-similarity. However, empirical relations between self-similarity, allometric relationships, and WBE’s length and radius scaling ratios remain ambiguous. Node-level average scaling ratios for the radius scaling ratio roughly support the theoretically predicted value even within individual tree crowns (Figure S4), agreeing with previous studies measuring this trait [33,37,39,86].

Selection for hydraulic efficiency in the node-level radius scaling ratio causes self-similarity in radial scaling [28]. This self-similarity also explains why cleaned trees have stable distributions of conductance exponents across branch pruning levels, illustrating the property of scale-invariance characteristic of self-similar structures (Figure S2). Conversely, within tree crowns, all average node-level measurements of branch length scaling, *γ*, deviate from the WBE prediction for space-filling in branch lengths (Figure S4). The space-filling length scaling ratio *γ* is therefore primarily responsible for deviations in calculated metabolic scaling exponents, relative to observed leaf area scaling exponents.

The correlation we measured between empirical leaf area scaling using allometric branching traits and theoretical metabolic scaling exponents (Equation 8) is weaker than expected if closed-form theory perfectly matched our empirically derived leaf area scaling. Instead, theoretical exponents exhibited low estimates of the metabolic scaling exponent (∼0.6), consistent with other work applying this approach [33,37,46]. We hypothesize that deviation from space-filling branching for ontogenetic or ecological reasons is the key cause of variation in *γ*, which may also be driving variation in leaf area scaling within individual tree crowns (Figure 4a, insets).

According to WBE, mass and leaf area scaling allometries should emerge from selection for space-filling branching. This is because both integrate variation in *γ* across tree crowns by combining length (and radius) scaling to represent network mass/volume. These curved scaling exponents (*c* and *θ*) depend on network depth, suggesting that self-similar assumptions for space-filling described by *γ* are violated due to variation in branch length scaling at different network depths. At the same time, allometric branching traits appear to integrate variation in node-level scaling ratios (as residual variation in regression lines) to capture approximate self-similarity and calculate scaling exponents consistent with WBE. Therefore, both scaling relationships are complex expressions of crown structure and function and may be subject to variability from many ecological and evolutionary processes.

A growing pool of evidence suggests that space-filling requirements do not tightly constrain plants since environmental and ontogenetic factors can influence branch length scaling [33,37–39,86,87]. Simple node-level estimates of space-filling may be biased by the presence of a few long branches, leading to extreme variability in branch length statistics [88,89]. These observations may indicate that space-filling is a more holistic measure of plant networks located at intermediate or crown scales, rather than at fine-scale branching junctions [40,46,90]. Length scaling is known to be more evolutionarily labile than the highly convergent area-preserving radial scaling in plant architectures [28]. The difficulty in precisely linking *γ* with WBE’s assumptions about space-filling and self-similarity in branching architecture hampers our understanding of how space-filling might evolve from adaptive pressures on tree architectures. It represents a critical theoretical and empirical gap in the WBE framework. Despite these biological realities, self-similarity remains a useful organizing principle for interpreting real-world variability, relative to simple theoretical predictions for tree architectural scaling encapsulated in allometric branching traits.

### Reconciling allometric variability with Metabolic Scaling Theory

Our results highlight the importance of accounting for variability in metabolic scaling due to multiple pressures on branching architecture evolution. The WBE network model provides a physiological foundation for Metabolic Scaling Theory (MST) and macroecology [91]. MST extends the allometric predictions of WBE to higher levels of organization, incorporating environmental variation and evolutionary history [24,92]. Therefore, WBE provides a biophysical foundation for making ecological predictions for individuals, populations and ecosystems within a single size-scaling framework [11,93]. Consequently, MST and WBE share many optimality assumptions and allometric predictions for optimal organismal functioning, while also permitting variation around the central quarter-power tendency for metabolic scaling.

The idea that variation in plant branching architecture may reflect differences in metabolic scaling and other allometric patterns in life history across species and environments has been advanced recently [29,90,94]. Variation around the optimal allometric predictions of MST has been observed across various dimensions of plant functioning over the past twenty years [25,31,32,95]. Empirical work has shown how tree-level allometries emerging from branch geometry, such as growth and crown traits, interact with environmental factors to produce variability in size-scaling [96–98]. Studies from forest plots have also shown that tree allometries exhibit wide variation relative to the optimal MST prediction [21–23].

A common alternative hypothesis is that variation in plant allometries (e.g. diameter growth rates) results from resource limitation and biotic interactions that restrict allocation to growth, regardless of the metabolic capacity of the branching network [96,99,100]. Our results indicate that accounting for variation in branching architecture may be necessary to clarify the extent to which ecological effects modify the allometric scaling of plant performance across species. If allometric patterns in branching architecture can be linked to proxies of metabolic performance *in situ* (such as growth rates, water flux, total leaf area), many inconsistencies with previous tests of MST may be resolved. Unifying these insights with the functional, organismal theory available in metabolic scaling theory could then significantly advance our ability to make regional and global-scale predictions of forest ecosystems [101,102]. TLS data from forest ecosystems promise to advance this aim by allowing precise measurements of tree architecture linked with other sources of ecological variation. Our work advances toward this goal by quantifying key axes of variation in tree architecture and best practices for applying them to TLS.

We used TLS scans to non-destructively measure metabolic scaling exponents in large trees. Our distributions of exponents control for various biases, refining the understanding of how WBE assumptions apply to tree architectural data. Revealing ¾ scaling from vascular network geometry using TLS data further validates the WBE model as a purely geometric approach to understanding metabolic scaling in macroscopic organisms. Predicting allometric relationships from vascular geometry is a key test of scaling theory and an aspiration for ecosystem science across vegetation types where allometric relationships are widely applied. Our work shows that TLS will continue to shed light on the diversity and function of branching architectures as monitoring techniques improve. By integrating our findings with the broader context of metabolic scaling, we provide a more nuanced understanding of the variability in allometric relationships. This approach not only tests the foundational hypotheses of the WBE model but also extends its applicability to diverse ecological settings. Our results underscore the importance of considering both theoretical predictions and empirical variability to advance the field of metabolic scaling theory and its applications in forest ecology.

#### Box 1

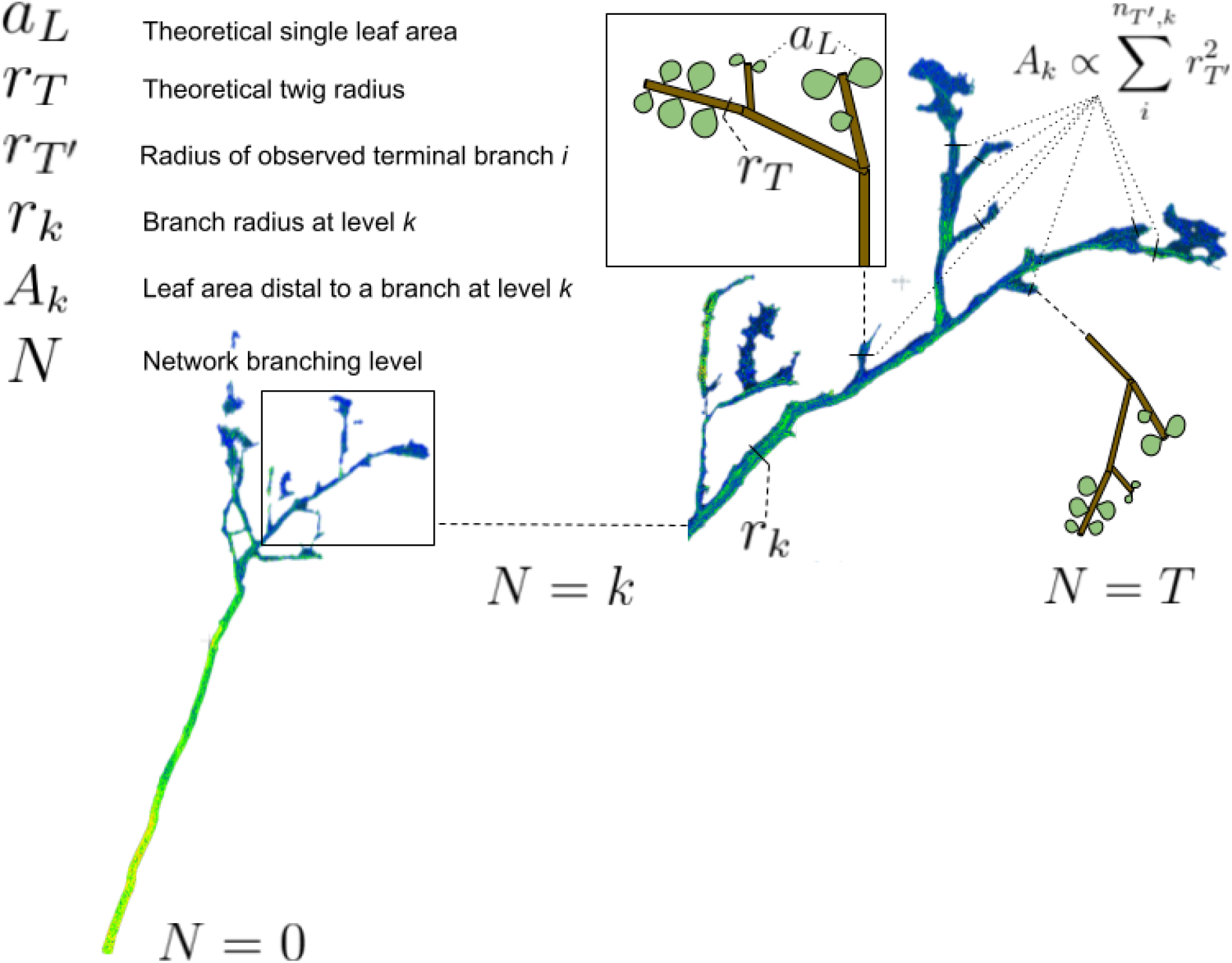

Box 1 : Pop-out cartoons of the twig scale show TLS terminals do not always reach twig and leaf depth T, instead terminating at a shallower terminal depth T’. Absence of twig scale could be a result of ontogeny, branch damage, or an artifact of TLS processing. We hypothesize that measured cross-sectional area of tenminal branches serves as an effective proxy of leaf area for analyzing network scal ing. This metric allows inferring the metabolic capacity of missing network segments and expllaining the scaling of the subtending proximal network. Symbols mapped to an actual scan of *Erythrophleum fordii*.

## Supplemental Information

### Figure S1

**Figure S1a:**
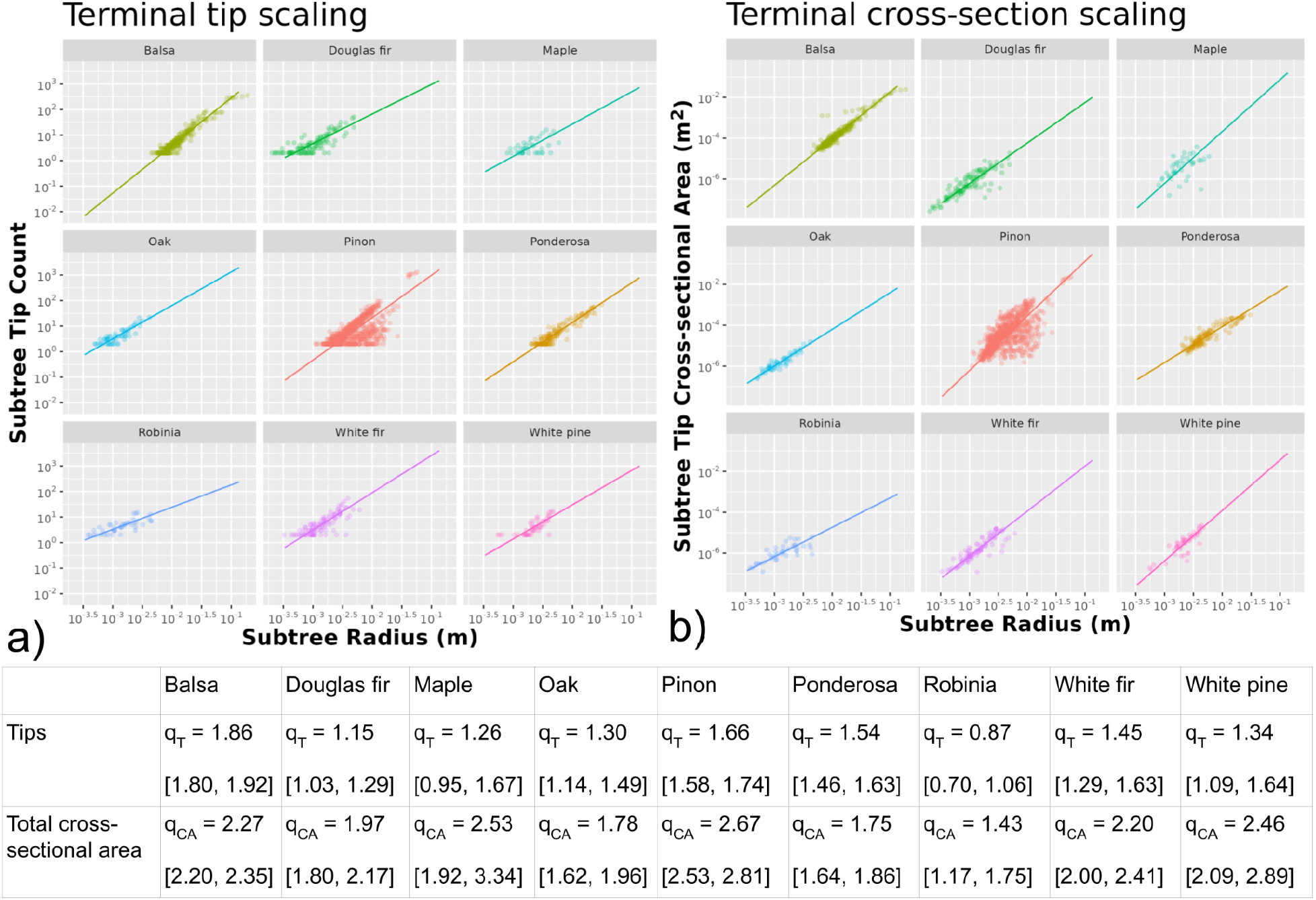
Terminal variability in manually measured plant networks match that of cleaned TLS scans. Two measures of the conductance scaling exponent *(q)* are given for multiple branching network samples in nine plant species. **a)** When counting terminal tips, radii vary by orders of magnitude, resulting in metabolic capacity estimates that bias toward shallow regressions. **b)** Instead measuring total distal cross-sectional area for a subtree accounts for variation in size that more accurately reflects WBE predictions. Table of predicted scaling slopes with 95% confidence intervals also provided. Interspecific terminal cross-sectional area scaling is not significantly different from the theoretical prediction: 2.11, [1.80, 2.43], p = 0.4174 in a one-sample t-test, while tip scaling predictions are significantly less (1.38, [1.15, 1.60], p < 0.001).

**Figure S1b:**
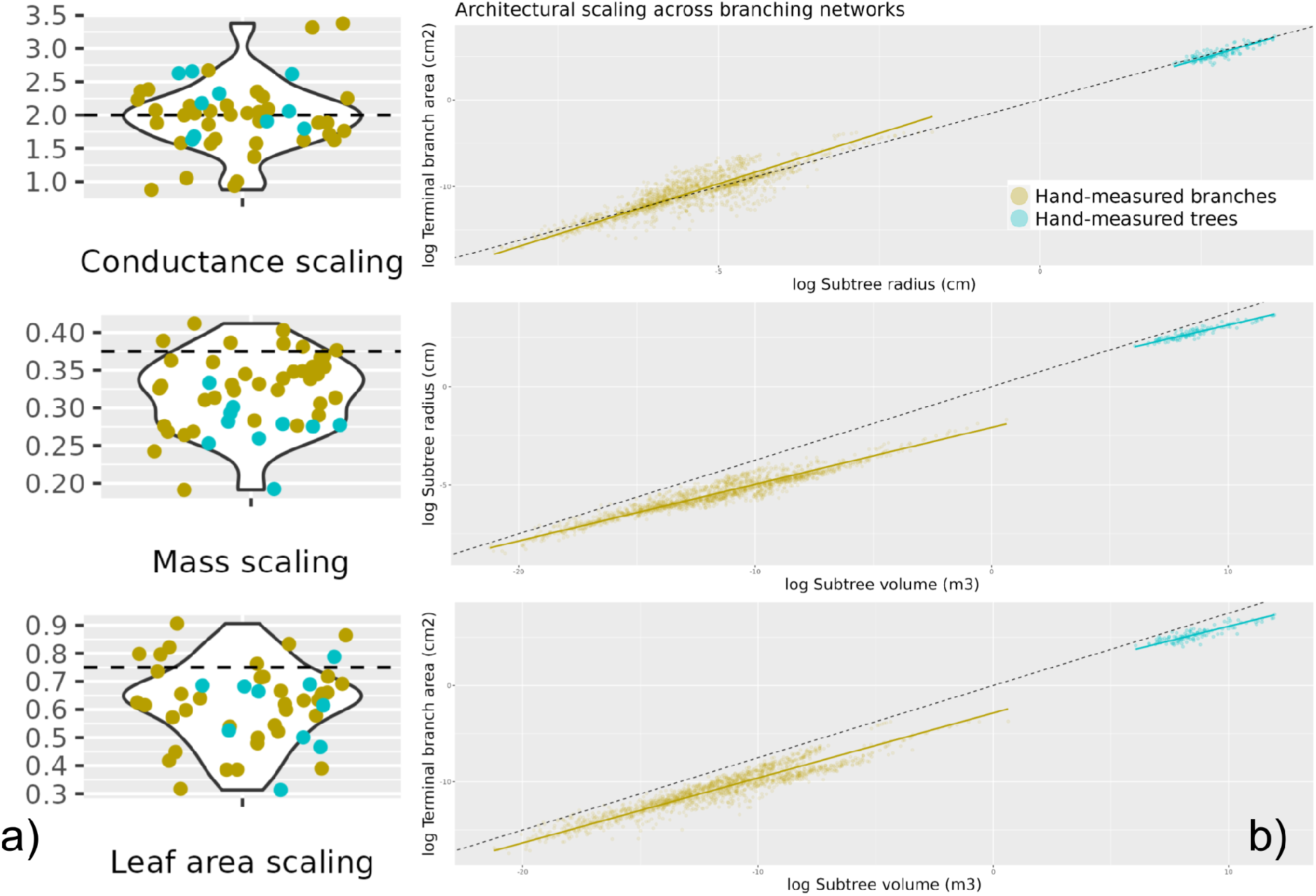
**a**) Scaling exponents from two sets of hand-measured plant data: one diverse set of branches from angiosperms and gymnosperms, and another dataset of felled tropical trees from Guyana. Points are scaling exponents for entire, individual networks (see Table S3). **a**) Scatterplot of subtrees across hand-measured plant network fragments. Points are individual subtrees across networks. Lines are SMA fits to the respective group of subtrees, dashed lines are theoretical predictions.

Results for hand-measured networks across an extremely broad range of sizes. We grouped plant vascular networks into two different size scales: hand-measured branches or trees and analyzed the same allometric patterns as in the main text. Distributions of exponents in panel **a)** are qualitatively similar to those observed for laser-scanned trees. In panel **b)** we show scatterplots and report interspecific regressions within and between the groups. The interspecific conductance exponent for hand-measured branches was 2.15 (2.13, 2.17), and for the Guyanese tree branches 1.88 (1.81, 1.96), and for both groups together 2.01 (2.00, 2.02). For the mass scaling exponent, hand-measured branches were 0.291 (0.289, 0.294) and Guyanese tree branches were 0.284 (0.272, 0.297), and across both groups 0.366 (0.364, 0.368). The metabolic leaf area scaling exponent for hand-measured branches was 0.628 (0.622, 0.634) and for the Guyanese tree branches was 0.537 (0.51, 0.56), across both groups giving 0.73 (0.734, 0.742). These results further emphasize the scale-dependence of these allometric relationships across diverse plant network geometries. Interspecific relationships are much closer to WBE predictions here, as in the TLS data we analyzed, indicating the importance of large-size-spectra and phylogenetic aggregations in picking out the convergent mechanisms hypothesized by WBE. Likewise, similar deviations in the finite-size affected allometries for individual networks (panel A) indicate that the same mechanisms influence variability at these size scales, indicating that unbiased TLS models reflect a ground truth for botanical branching data. Therefore, although not analyzed here, asymptotic patterns of variation appear across scales in manually measured morphological data, which support all our work’s main findings and validate the cleaning process as a prerequisite for TLS studies of branching allometry/architecture.

### Figure S2

Here, we include an approach to pruning small branches in QSMs. This analysis is motivated by a key consideration: how the error in small branch geometry from TLS data affects scaling estimates of allometric relationships in tree branch networks. We use scaling exponents computed across pruning treatments to illustrate the pitfalls of studying branching architecture in noisy TLS scans of tree crowns.

Pruning appears to degrade estimates of scaling in cleaned trees. Based solely on comparison with distributions of exponents for unbiased trees, pruning analyses may still be helpful if cleaning is not an option. There is an indication that an intermediate pruning level (50-75% branch removal) for unprocessed trees, particularly foliated scans, excludes small, overfit cylinders while retaining a significant fraction of network scaling.

#### Cylinder pruning analyses

As an additional control on the quality of structural models, we investigated the removal of small branches which may bias estimates of network scaling. We conducted cylinder pruning analyses on QSMs using quantiles on branch radii within a network. In each pruning analysis, we removed all branches below a given size threshold. Connect distal branches were removed if an interior branch (proximal to the trunk) were removed. Allometries were then re-computed for each pruned tree. Five percentiles were included: (0%, 25%, 50%, 75%, 90%). We repeated this analysis with absolute branch radius cutoffs in meters (0.0025, 0.005, 0.01, 0.025, 0.05) and branch orders (1-5) with similar results. Quantiles are most appropriate for analyzing trees of different sizes where the minimum and maximum branch sizes can vary between networks. An example of this protocol is presented in Figure S2a. Statistical details for pruning treatments are available in Table S1.

#### Branch pruning effects on network scaling

When analyzing tree cylinder models derived from TLS, terminal branches exhibit high error and the true terminal size scale is usually absent from cylinder models. This error may influence estimates of the scaling exponent. Selectively pruning the distal branches of the scanned network affected by error is a potential solution. While scaling may vary in the portions missing from the network, pruning hypothesizes that approximate self-similarity of branch dimensions allows us to recover network scaling relationships, and we briefly prove that here.

Pruning is defined as removing all branch segments up to a level *T′* such that the new network depth *k* < *T′*< *T*. The first key result in this context is that the scaling in Equation 5 is unchanged for all network levels. This only involves modifying the expression in Equation 4b or Equation 5 to index a new network depth *T′* resulting from pruning, just as in our discussion of distal leaf area scaling. Most importantly, this change appears in Equation 4b as the constant of proportionality between leaf area and terminal branch cross-sectional area,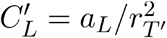 Therefore, when branch radii are self-similar in a network, pruning will only shift the intercept, rather than the exponent of leaf area scaling relationships.

Self-similarity also preserves the expression for stem-mass allometry when branches are pruned. However, we expect branch removal to reduce the curvature in Equations 4 and 5, allowing straight-line fits to better estimate the metabolic scaling exponent *θ* (and the mass allometry *c*). This is because pruning removes subtrees where *k* ≈ *T* such that *A*_*k*_ ∝*V*_*k*_.

Pruning analyses seem to affect allometries with asymptotic components of variation more than the conductance allometry, but bias makes this unclear.

**Figure S2a:**
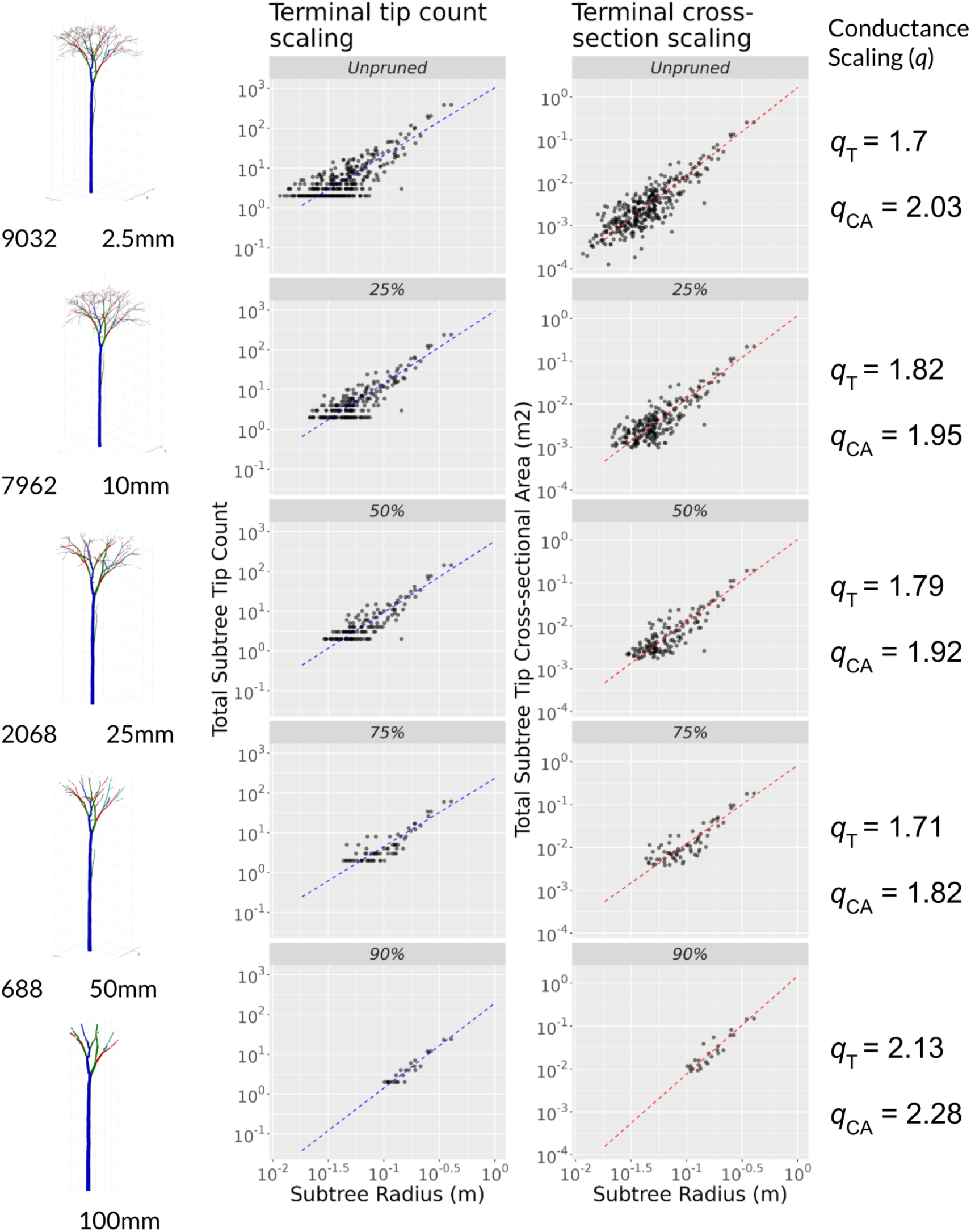
*Left)* Example of the branch pruning process across five quantiles of branch radius within an unbiased (cleaned) *Cylicodiscus gabunensis* individual. Total branch number and minimum branch size are listed up to the 90th percentile. Each round of pruning produces a new set of terminal tips for analyzing network scaling while decreasing the total size of the network analyzed. *Center)* Regressions on two subtree distal leaf area estimates: Integer tip count versus terminal cross-sectional area. Dashed lines are bivariate standardized major-axis fits to Eq. 5. Note that cross-sectional area re-scales the vertical axis by varying the terminal dimension with size. *Right)* Fitted values of the conductance allometry for terminal tips (q_T_) and terminal cross-sectional area (q_CA_) regressions.

**Figure S2b:**
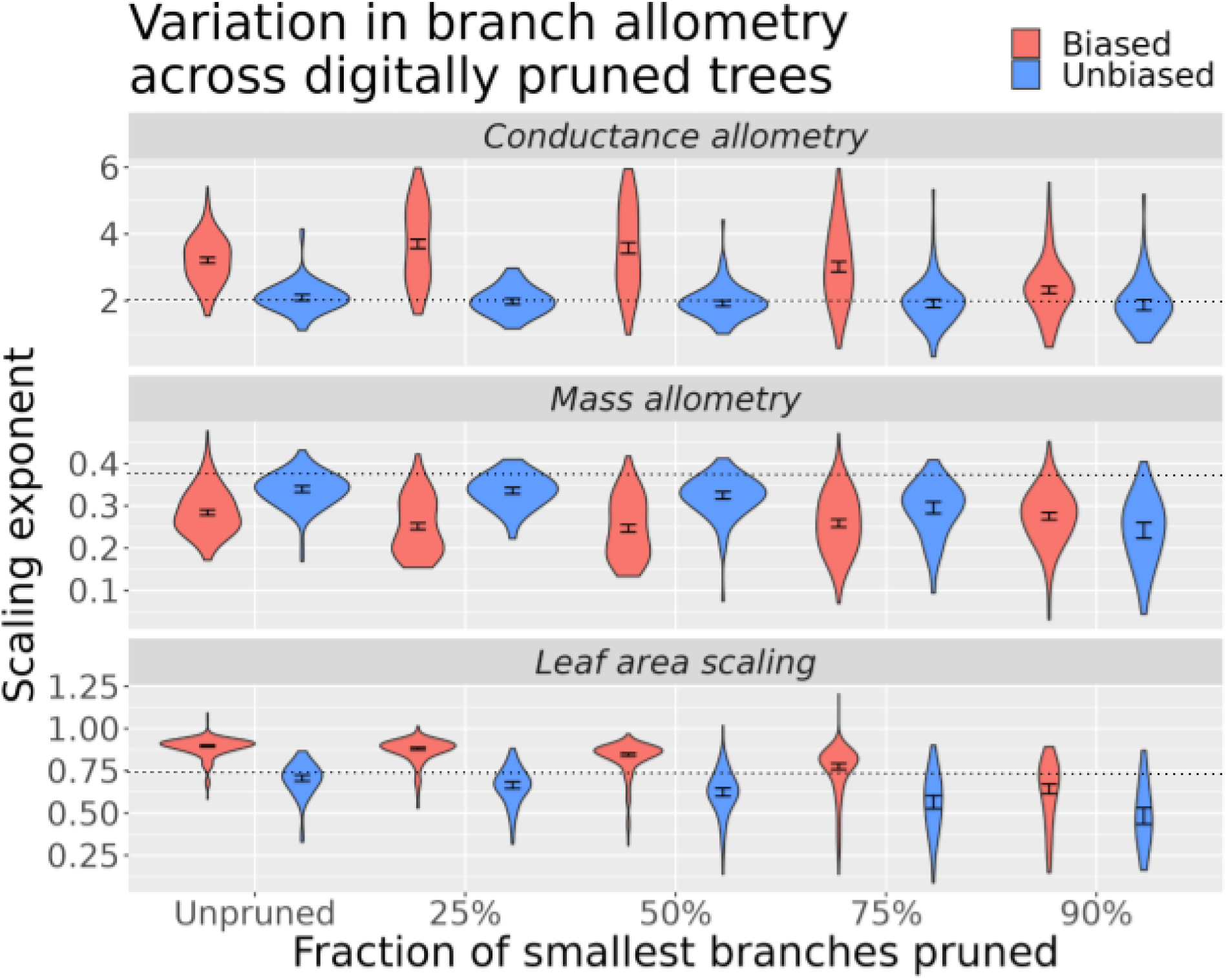
Distributions of empirical scaling exponents across quantile thresholds. Distributions are given for two processing methods: raw cylinder models, and cleaned cylinder models with defoliation and editing. Dashed line is predicted optimum exponent from metabolic scaling theory for each architectural allometry. 95% confidence intervals from Wilcoxon tests are shown for each distribution.

Distributions of scaling exponents for three architectural allometries (conductance scaling *q*, stem mass allometry *c*, and leaf area scaling *θ*) across quantile prunings are shown in Figure S1. Pruning branches tends to increase the overall variability in scaling observed across individual trees, likely due to reducing the number of branches sampled. Pruning can shift scaling estimates closer to theoretical predictions in raw structural models, but has variable effects on cleaned trees. 95% confidence intervals derived from one-sample Wilcoxon tests are significantly different from theoretical expectations for all distributions except for conductance allometries in cleaned trees, which match theoretical expectations across all pruning levels.

Cleaned and raw trees systematically varied in their statistical properties across quantiles, presented in Table S1. Cutoffs around 25-50% tended to exclude branches up to about 1-3cm radius, with 90% cutoffs excluding ∼5-10cm branches. Pruning analyses consistently reduce branch number, causing the number of whole trees eligible for regression analysis to drop due to sample size constraints. This is because the total network branch number drops below 7 for some individuals (3 subtrees), after which regressions cannot be conducted (n < 3). Prior to this, small numbers of branches will still result in highly variable regression slopes. Importantly, cleaned trees tended to exhibit larger minimum branch sizes, as the cleaning process inherently removes smaller branches poorly captured by TLS workflows or erroneously introduced in QSM-fitting, particularly segments <1cm.

### S2 Table - Tree statistics across quantile pruning thresholds

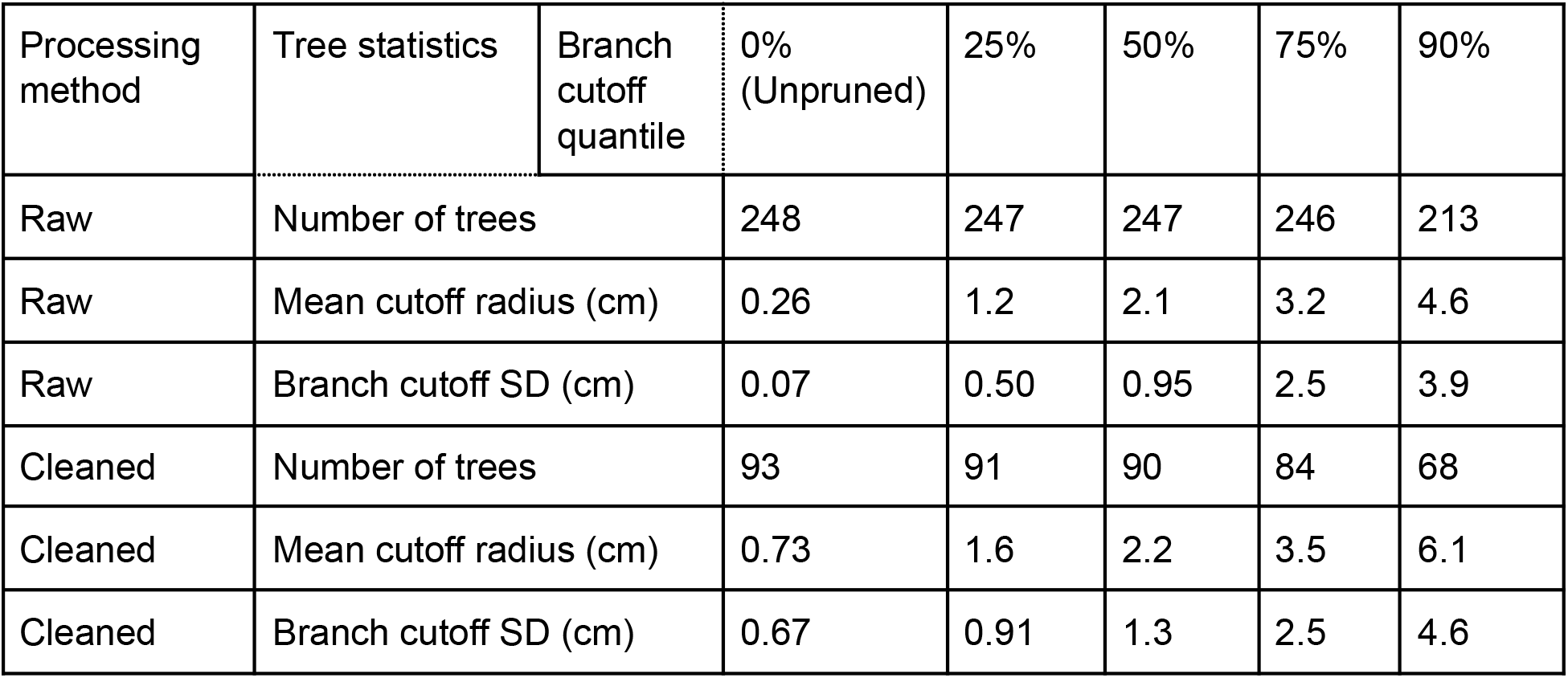

### Figure S3

**Figure S3:**
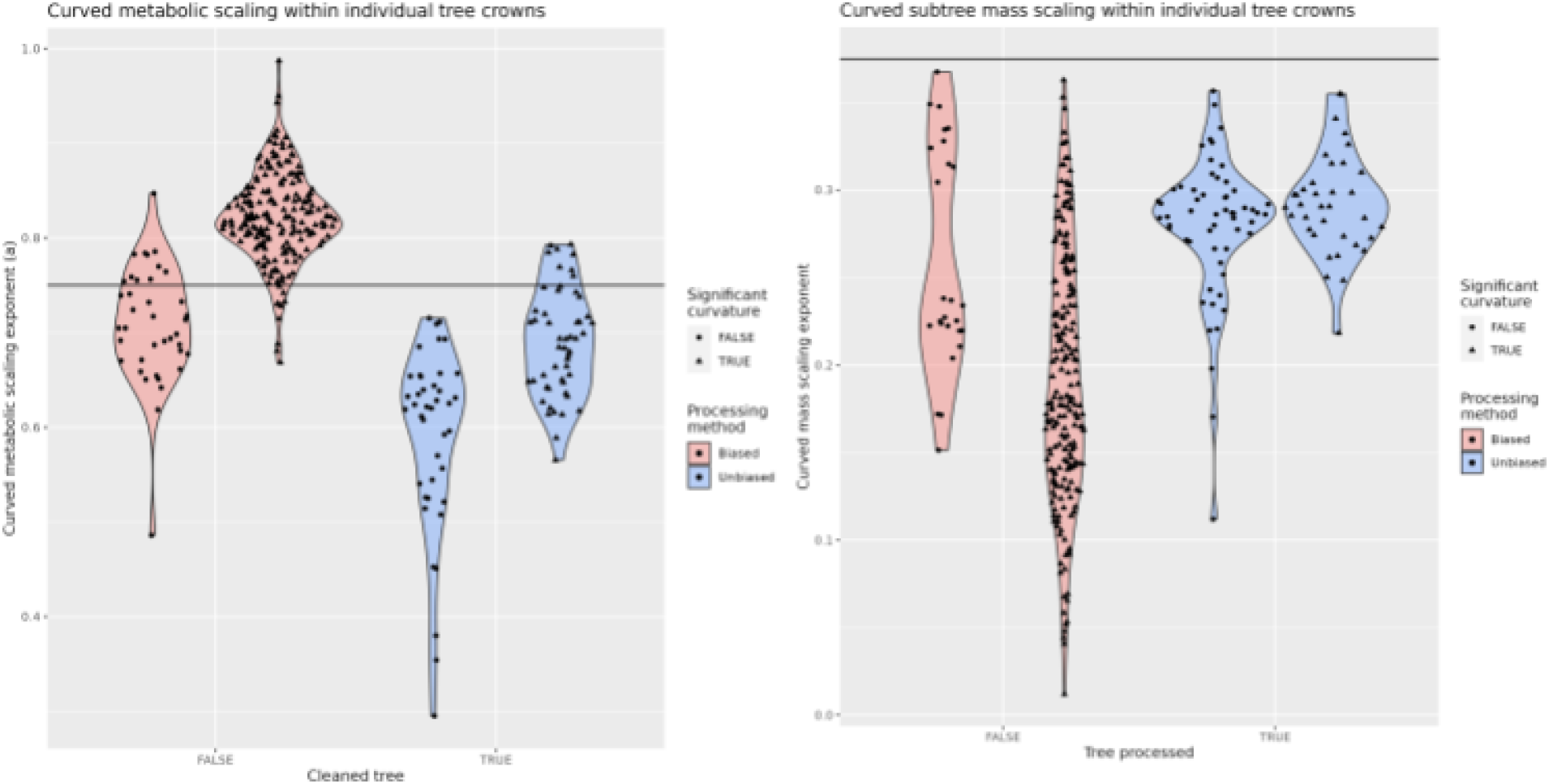
Results of curved regression experiments within individual tree crowns. Horizontal lines are the theoretiical expectations for each exponent. Overall patterns across processing methods are similar to straight-line regressions, as are distributions of scaling exponents.

Figure S3 is the result of fitting curved functions within individual tree crowns, for all individuals in the dataset. To maximize the chance of detecting curvature, we present only unpruned networks. Points show the value of the parameter *a*, and shapes dictate whether the curvature parameter *b* was statistically significant in the regression. Estimates of scaling are generally comparable to those obtained from RMA regressions in the main body, most importantly the difference between cleaned and raw trees. In either case, curvature is not significant in a large minority of trees.

Intraspecific and intraindividual scaling exponents do not reflect theoretical predictions even when accounting for finite size effects with curvature. As indicated in the main text, only aggregated interspecific analyses with large branch size variation reflect WBE predictions.

Theoretically, curvature should become less apparent as more distal branches are pruned. In practice, non-linear fits become highly unstable as the sample size within a crown is decreased. Straight-line bivariate fits are generally recommended as being more robust to variability present in QSMs derived from TLS data, although future improvements in QSM quality, as well as intraspecific data for representing ontogeny might clarify curved metabolic scaling in trees in future research.

### Figure S4

**Figure S4:**
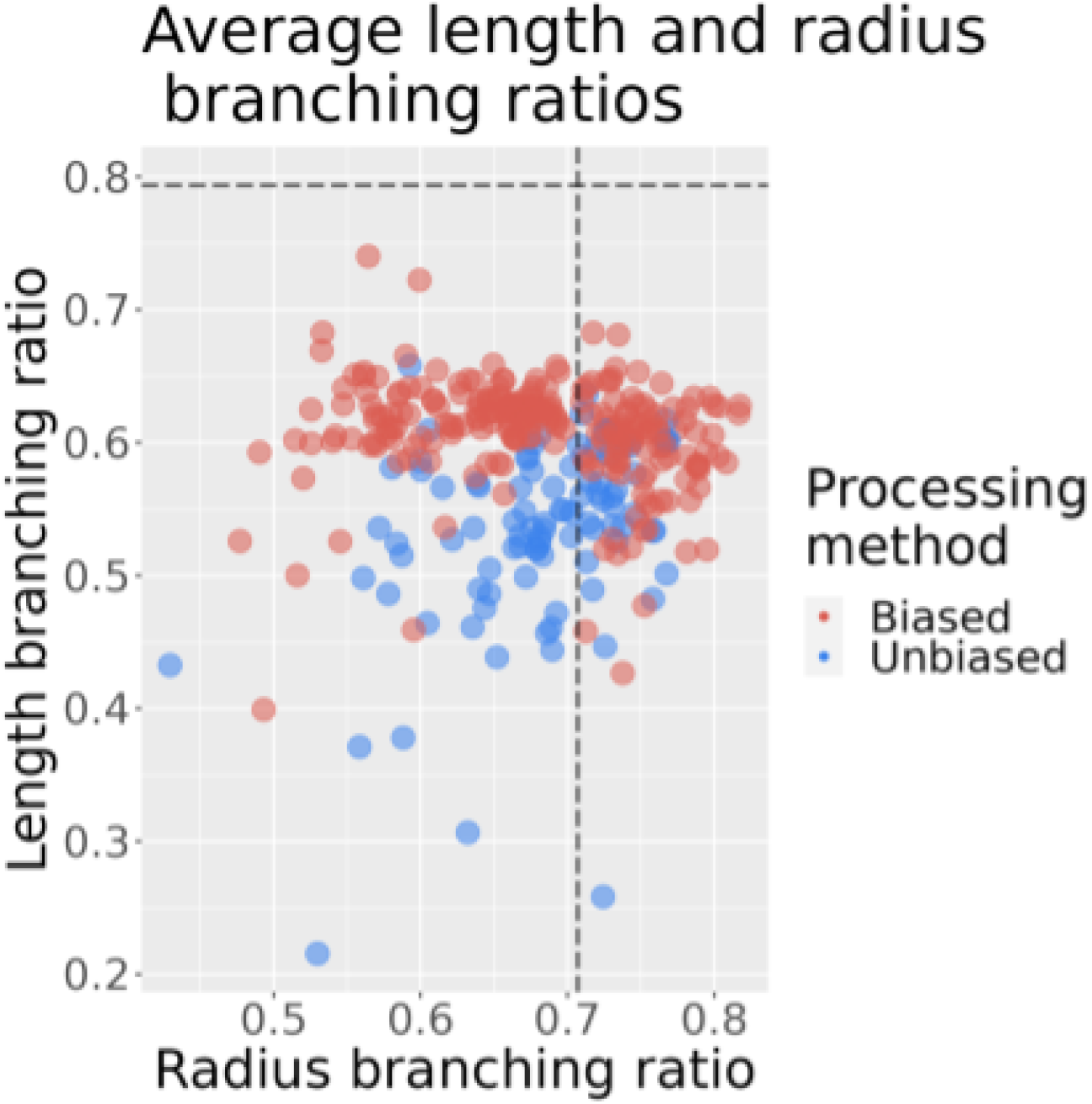
The radius Scalling ratio (chi1d branch radius divided by parent radius) is shown on the horizontal axis, and the length scaling ratio (child branch length divided by parent. length) is shown on the vertical axis. Data. points are individual tree averages. Horizontal and vertical lines represent the optimal values as predicted by WBE theory.

Branch scaling ratios for whole trees are shown in Figure S4. These are averages of ratios measured at each node within a tree. Radius scaling ratios were roughly predicted by WBE theory, while length scaling ratios diverged from the theoretically optimal value. Average length scaling ratios were apparently lower in cleaned trees, perhaps due to the removal of slender twigs and branches at the terminal ends of the network.

### QSM Supplement

The core architectural relationships we studied also reveal a key signature of allometric bias in TLS-derived structural models. Scaling exponents from unbiased data are clustered relatively close to the predictions of WBE, and we observed that this range of exponents matches that of manually measured plant networks, providing some ground truth to our scaling estimates for TLS models (Figure S1b). Conversely, biased tree data exhibited extreme variation in both exponents underlying metabolic scaling. Analyzing metabolic scaling with regression-based allometric branching traits relies on estimating terminal branch geometry from TLS data. Since the conductance exponent is calculated using the smallest, terminal cylinders in a given network, smaller branch radii are a key determinant of QSM quality for studies of branching allometry.

Overfitting cylinders clearly cause bias throughout the branching network, and cleaning and validating trees manually removes. Essentially, the cleaning process reduces the number of fitted cylinders by an order of magnitude, mainly by human-corrected algorithmic defoliation, and reduces skew in the size distributions of branches (Figure S6). The strong signal of bias explains the drastic variability in allometric exponents in the broader dataset, contra WBE predictions, and confounds much of any residual natural signal of variation. Distributions of exponents from the broader dataset of TLS trees are artifacts of the regression procedure paired with systematic bias in the underlying cylinder geometry.

Therefore, we have illustrated significant differences in the focal allometric exponents that reflect the broader pattern in global TLS data compiled. The Figure S5 makes clear how biased tree models produce erroneous estimates of allometric scaling. Biased trees have a consistent pattern in branch size distribution that explains deviations in allometric predictions.

We relied on the TreeQSM (v2.4.1) software suite and leveraged the recommendations available in the TreeQSM manual, as well as recent work on TreeQSM parameter optimization, particularly (Gonzalez de Tanago et al 2018) and (Demol et al 2022). From these works, we identified a range of values for the key parameters PatchDiam2Min and PatchDiam2Max that could fit trees well across a range of scanning conditions, instruments and data types.

PatchDiam2Max = [0.07 0.1 0.15 0.25]

PatchDiam2Min = [0.025 0.05 0.075 0.1]

PatchDiam2Max mostly regulates fitting of the trunk and large relay axes, and is relatively robust to variation in scan quality. Optimal QSMs tended to minimize this parameter closer to the 7cm value. PatchDiam2Min mostly governs fitting smaller, sensitive terminal branches and twigs. Because it tended to overfit, we set the PatchDiam2Min beginning at the recommended 2.5cm and only allowed larger patch sets. This is likely not appropriate for the highest-quality scan data (for example, our highest quality scans of *Ulmus* individuals from Alberta, Canada were better fit by a PatchDiam2Min of ∼0.005-0.01). Therefore we consider our QSMs to be a conservative set of fits tending to exclude erroneous terminal branches. However, we retained the default minimum branch size (2.5mm) in TreeQSM, which is below the scanning resolution of all of our data sources. The accumulation of overfit cylinders at this minimum size is also apparent in scatterplots and frequency distributions of branch sizes for biased trees. Constraining this parameter to more realistic minimum branch sizes is recommended when fitting to tree clouds that have not been defoliated and manually corrected.

Most trees tended to match the optimal parameters identified in the studies above, particularly an optimal PatchDiam2Min value around ∼2.5cm, especially in dense and/or foliated point clouds. However, this core parameter will tend to overfit trees, creating ‘reticulated’ fits on branch surfaces and erroneous small cylinders (roughly less than 1cm in diameter), especially within foliated and occluded scans (see screen captures below).

We have found that the proper selection of optimization criteria within TreeQSM’s optimization routine can lead to robust fits across heterogeneous scanning conditions, foliation states, and point cloud sparseness. These settings typically only need to be identified once per scanning dataset/field conditions. In particular, we identified three error criteria that are suited to different datasets, listed as ‘optim’ in Table S3:

‘all_mean_dis’: the default error criterion in TreeQSM will tend to result in overfitting and the selection of QSMs with many erroneous cylinders. However, this criterion frequently accurately identifies the QSMs that fit intermediate and fine-scale branches. This criterion most motivates branch pruning/removal prior to scaling analysis due to its tendency to fit small (millimeter-scale) branches to noise. This becomes particularly important when fitting foliated scans - cylinder removal is vital, as many cylinders below 10mm are effectively fit to leaf points. At the same time, other optimization criteria typically cannot handle foliated trees, instead fitting extremely large (5-10cm) branches to leaf points.

‘all_mean_surf’: this alternative criterion optimizes the distance of points from the surfaces of cylinders, which is perhaps more appropriate for studies of branch architecture focused on the conductive capacity of branches and the dimensions of cylinders (rather than the volume per se, as in studies of volumetric biomass estimation). More importantly, this criterion tends to penalize the production of many small and reticulated cylinders from noise in the distal ends of tree networks, resulting from other known sources of error in tree scans (coregistration error, occlusion error). As a more conservative criterion than ‘all_mean_dis’, it may fail to fit cylinders to branches present within a tree scan.

‘all_min_surf’: rather than minimizing the mean distance from cylinder surfaces, this minimizes the minimum distance across QSMs. This extremely conservative criterion is most appropriate for fitting very noisy and minimally branched individuals in specific cases. Realistically, such trees should be manually cleaned according to the methods described in this paper.

We identified ideal optimization criteria for all trees in the dataset (See Table S3). Typically, similar species and scanning conditions will lead to a particular criterion being suitable for a specific dataset. Still, some datasets require consideration of trees on a case-by-case basis. When optimization criteria produce similar results, a secondary criterion is to choose the fit that minimizes the total number of cylinders, which tends to imply fewer, larger cylinders as more restrictive optimization criteria are chosen. In turn, this optimizes for modeling the well-constrained volume of larger branches in the network hierarchy. The embedded figure depicts this trade-off in branch number and branch volume for the two main optimization criteria used. A one-to-one line is visible in the data where the two criteria are interchangeable for many networks. We typically used ‘all_mean_surf’ to break ties when fits were similar. Future work may formalize this heuristic when fully automating QSM fitting in scaling analyses.

**Figure S5:**
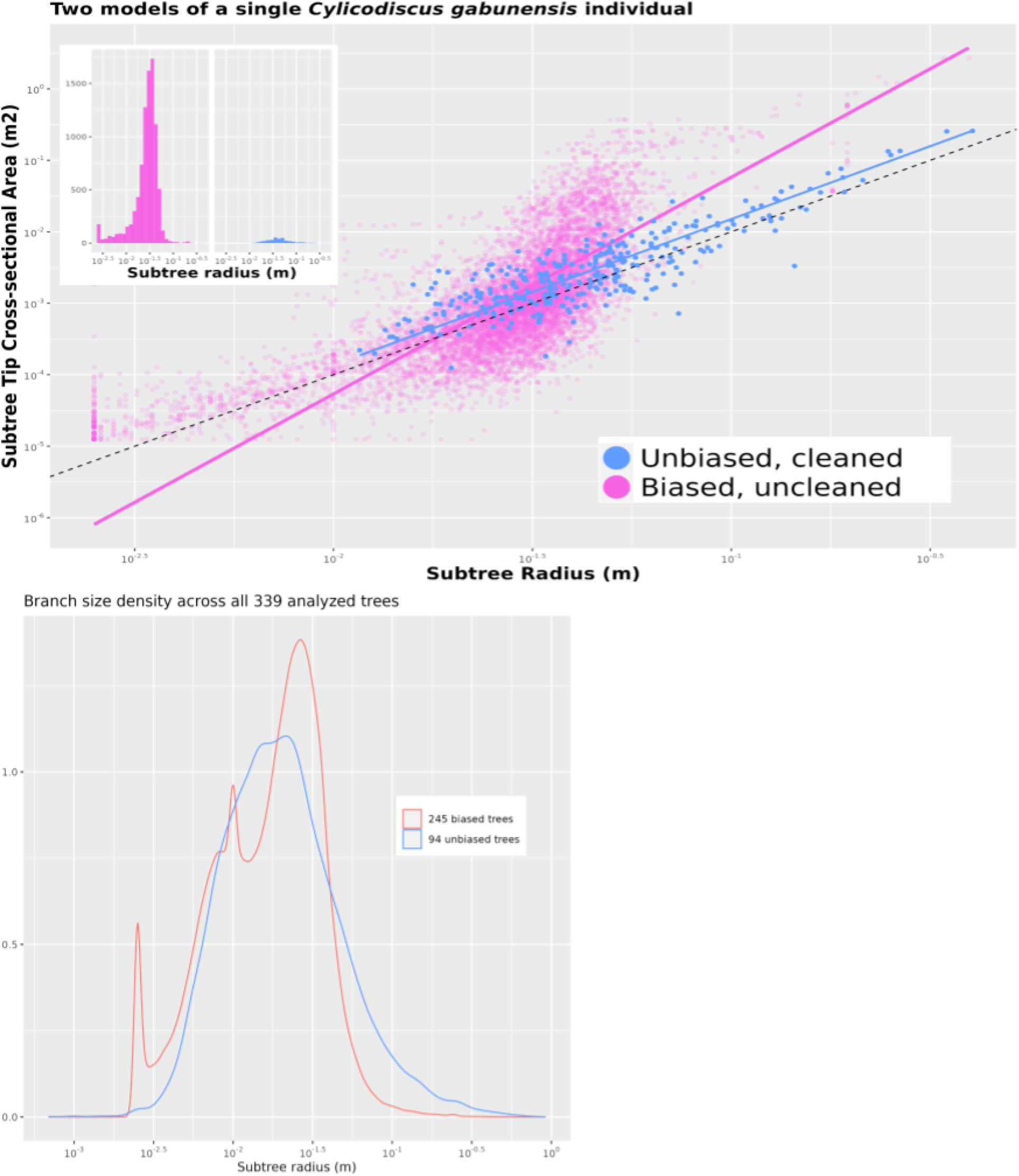
**a)** Scatterplot of all subtrees for two models of a *Cylicodiscus* gabunensis individual. Magenta points drawn from an unprocessed structural model and blue points are from another model subjected to cleaning, removing bias. Unbiased trees greatly reduce the number of fitted branch cylinders and generally restrict branch geometry to the centimeter scale and above. **Inset:** In the unprocessed *C. gabunensis*, an order of magnitude more cylinders are fit, extending to millimeter scales. Density plots of subtree branch radius distributions show clear signals of overfitting through spikes in density distributions around 1 millimeter and 1 centimeter, reflecting different processing techniques across the broader global dataset of biased tree models

**Figure S6:**
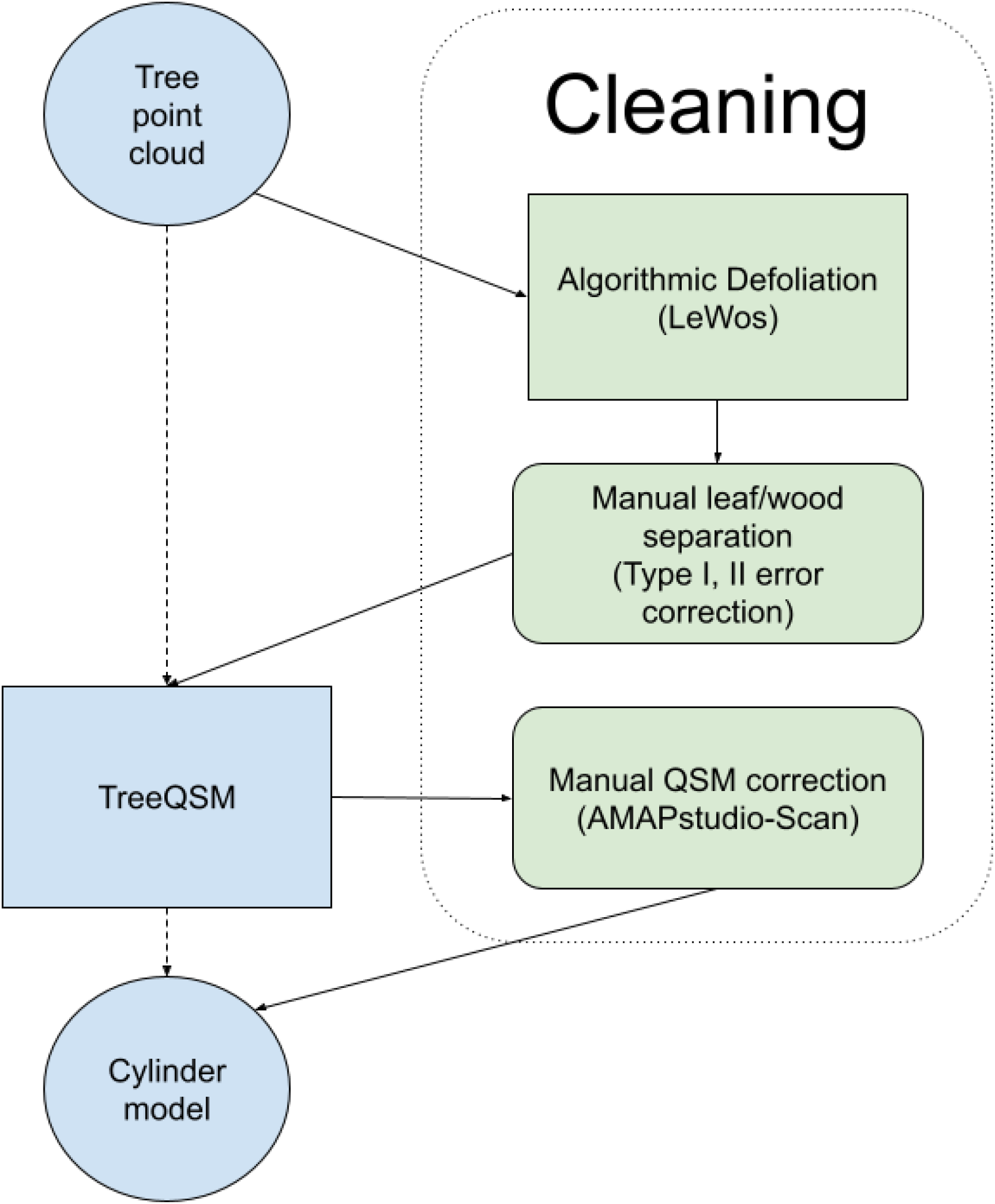
Simplified flowchart for cleaning structural models, from processed input cloud to final cylinder model. For our analysis, the dashed line represents ‘raw’ trees, while solid lines represent the cleaning process used for two different datasets

Cleaning is a straightforward process for reducing errors in tree processing, but it is not easily reproducible due to the need for manual intervention. In this work, the authors responsible for analysis did not participate in tree cleaning, and cleaning was conducted prior to conceptualization of this work. We submit that the potential for confirmation bias and manipulation of data is minimal in these cases.

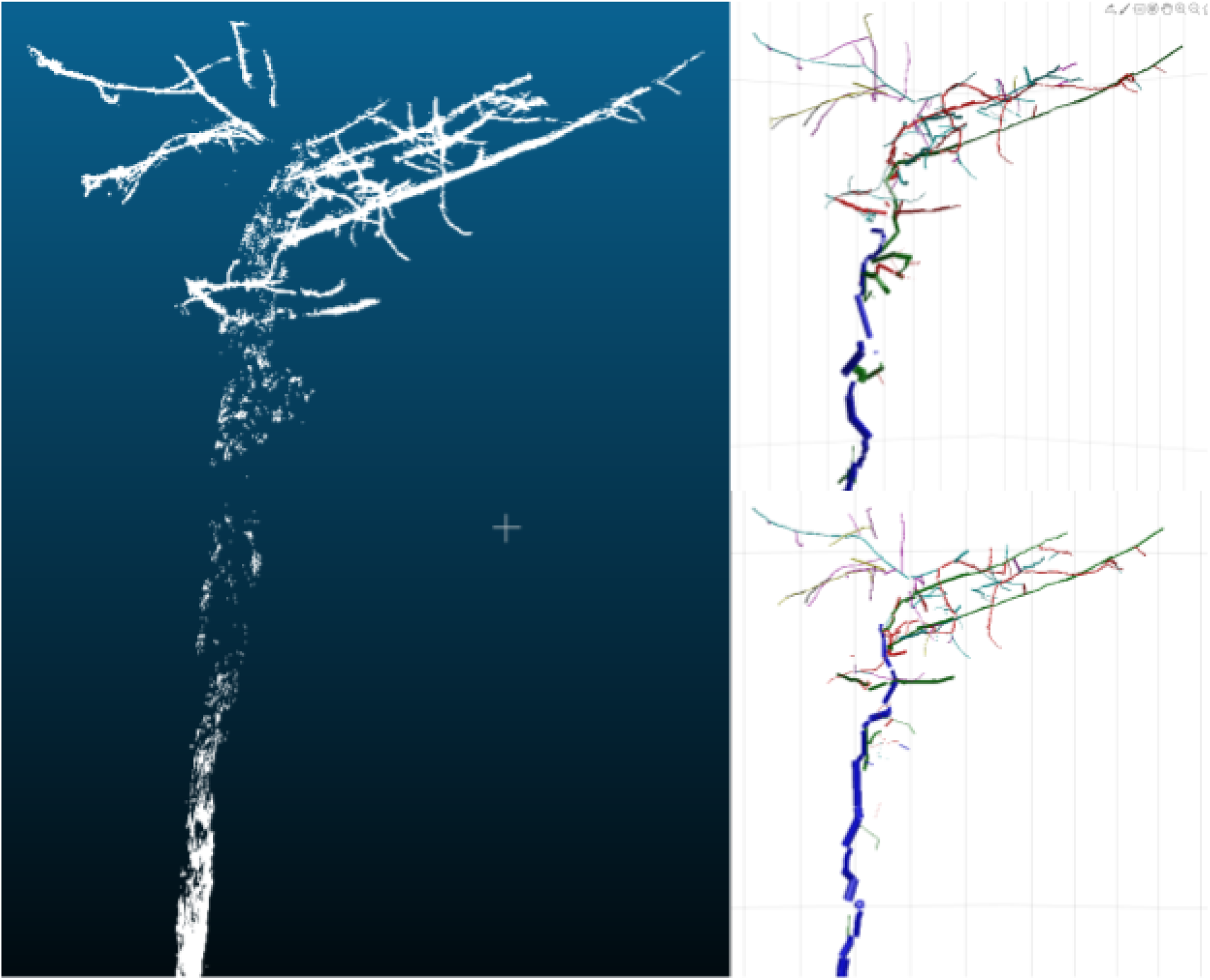

A noisy and occlluded *Pycnanthus angolensis* with g1aps in fine branching. At top right,, optimization ‘all_mean_dis’fits erroneous. cylinders to the occluded main axis andfine branches. At bottom right, ‘all_mean_surf’ recovers the main axis and more conservatively fits fine branches in the miinimal crown availablle.

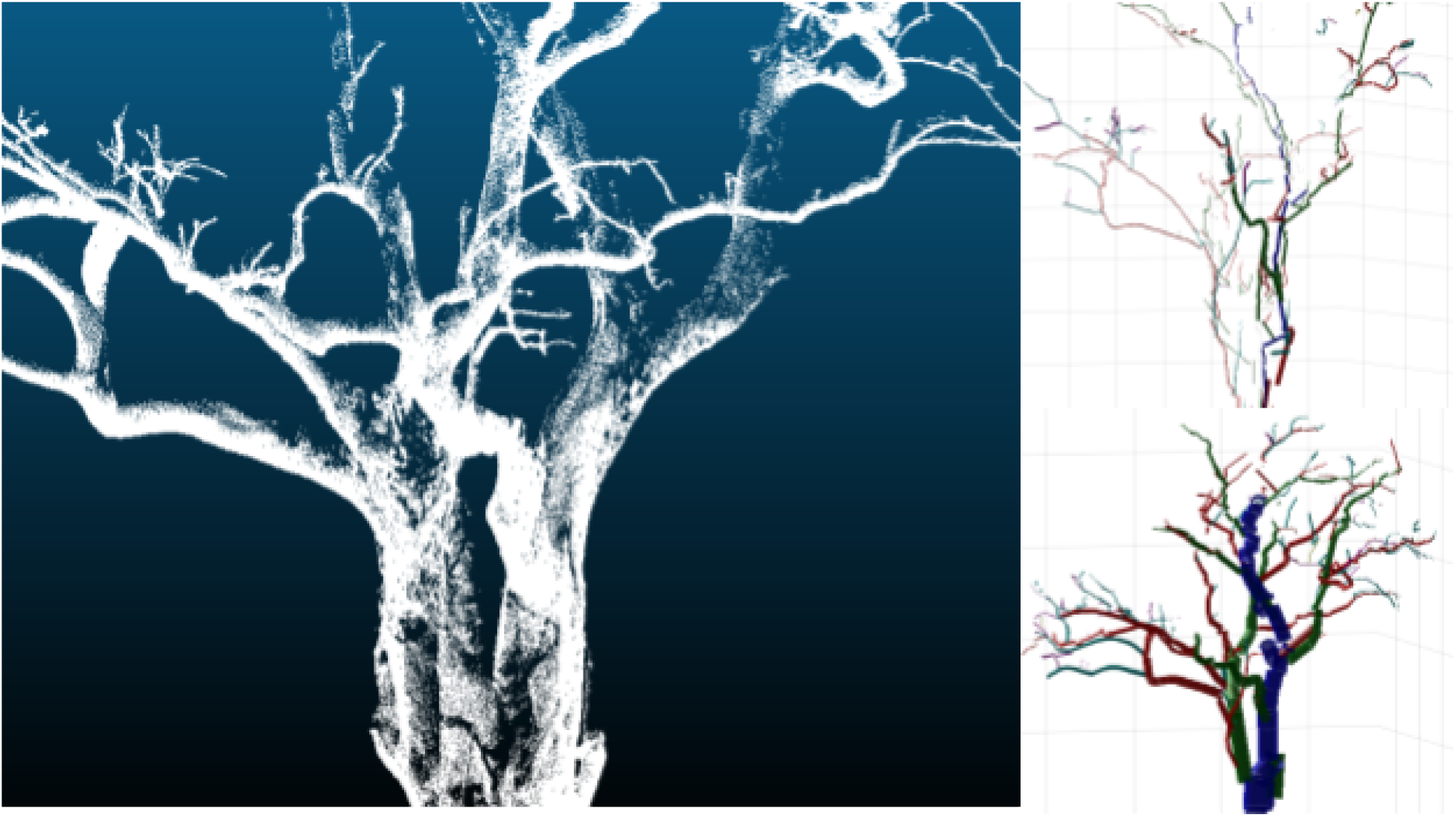

Variation in the effiicacy of optimization criteria on an occluded scan of *Triplochiton sclaroxylon*. At top right, the criterion ‘all_mean_surf’ and ‘all_mean_dis’ fit cylinders to the contours of occluded branches. The more, conservative criterion ‘all_min_surf’ optimizes mini mum distances to cylinder surfaces and fits volume more property.

**Table S2:** Data sources with the geographical location, number of trees, foliation status of the trees, cleaning status, scanner used, and reference in the literature

| Code | Location | Trees | Cleaned | Scanned<br>Leaf-on | Instrument | Resolution<br>(mm) | Source |
| --- | --- | --- | --- | --- | --- | --- | --- |
| HACK1 | Germany | 24 | N | N | Z+F<br>IMAGER<br>5010 | 4 | Hackenberg et al<br>2014 [103] |
| HACK2 | China | 24 | N | Y | Z+F<br>IMAGER<br>5010 | 4 | Hackenberg et al<br>2015 [104] |
| HACK3 | Germany | 12 | N | N | Z+F<br>IMAGER<br>5010 | 4 | Hackenberg et al<br>2015 [104] |
| TECTONA | Ecuador | 3 | N | Y | FARO Focus<br>3D X-330 | 2 | Aguilar et al 2019<br>[105] |
| OXFORD | United<br>Kingdom | 15 | N | N | Riegl VZ-400<br>3D | 5 | Author data |
| GONZ1 | Guyana | 10 | N | Y | Riegl VZ-400<br>3D | 5 | Gonzalez de Tanago<br>et al 2018 [63] |
| GONZ2 | Indonesia | 10 | N | Y | Riegl VZ-400<br>3D | 5 | Gonzalez de Tanago<br>et al 2018 |
| GONZ3 | Peru | 9 | N | Y | Riegl VZ-400<br>3D | 5 | Gonzalez de Tanago<br>et al 2018 |
| ALBERTA1 | Costa Rica | 19 | N | Y | Riegl<br>VZ-400i | 5 | Guzman-Quesada et<br>al 2020 [106] |
| ALBERTA2 | Canada | 15 | N | N | Riegl<br>VZ-400i | 5 | Guzman-Quesada et<br>al 2020 |
| CALD | Australia | 65 | N | Y | Riegl VZ-400<br>3D | 5 | Calders et al 2015<br>[107] |
| ZOFIN | Czechia | 43 | N | Y | Leica C10<br>Scanstation | 5 | Trochta et al 2017<br>[108] |
| MOMO | Cameroon | 61 | Y | Y | Leica C10<br>Scanstation | 5 | Momo et al 2017 [66] |
| AMAP | Guiana | 33 | Y | Y | Leica C10<br>Scanstation | 5 | Author data |

**Table S3:**
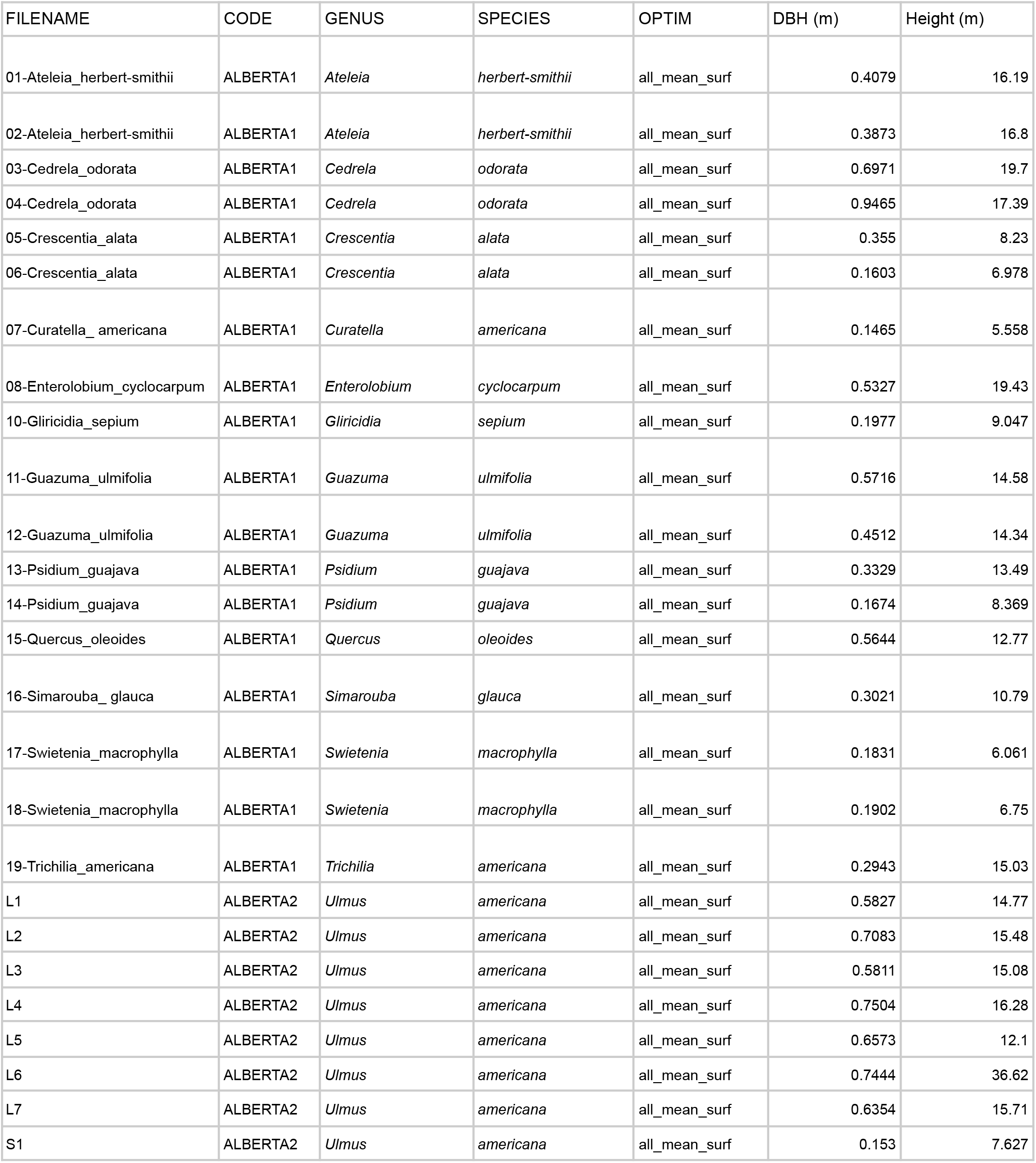
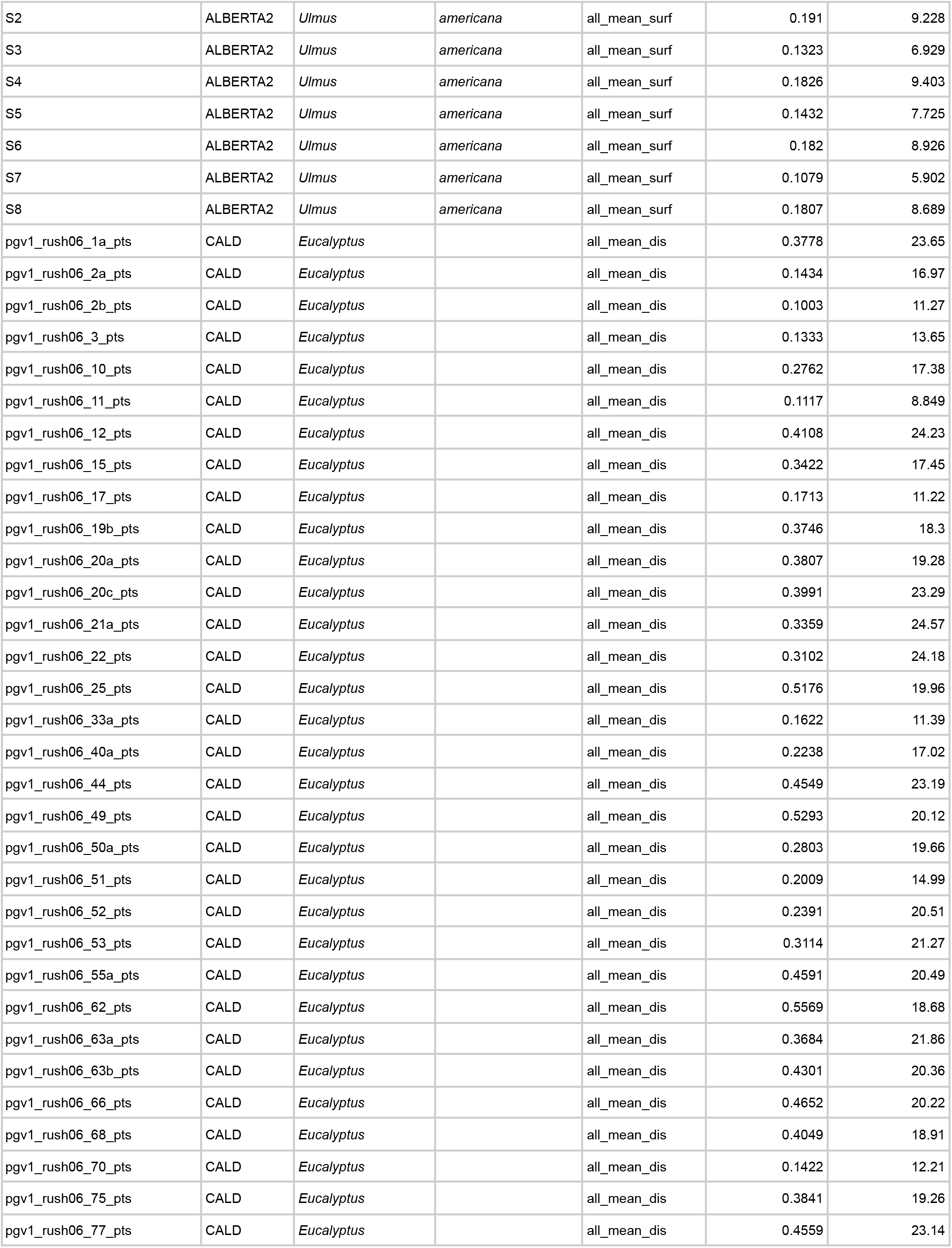
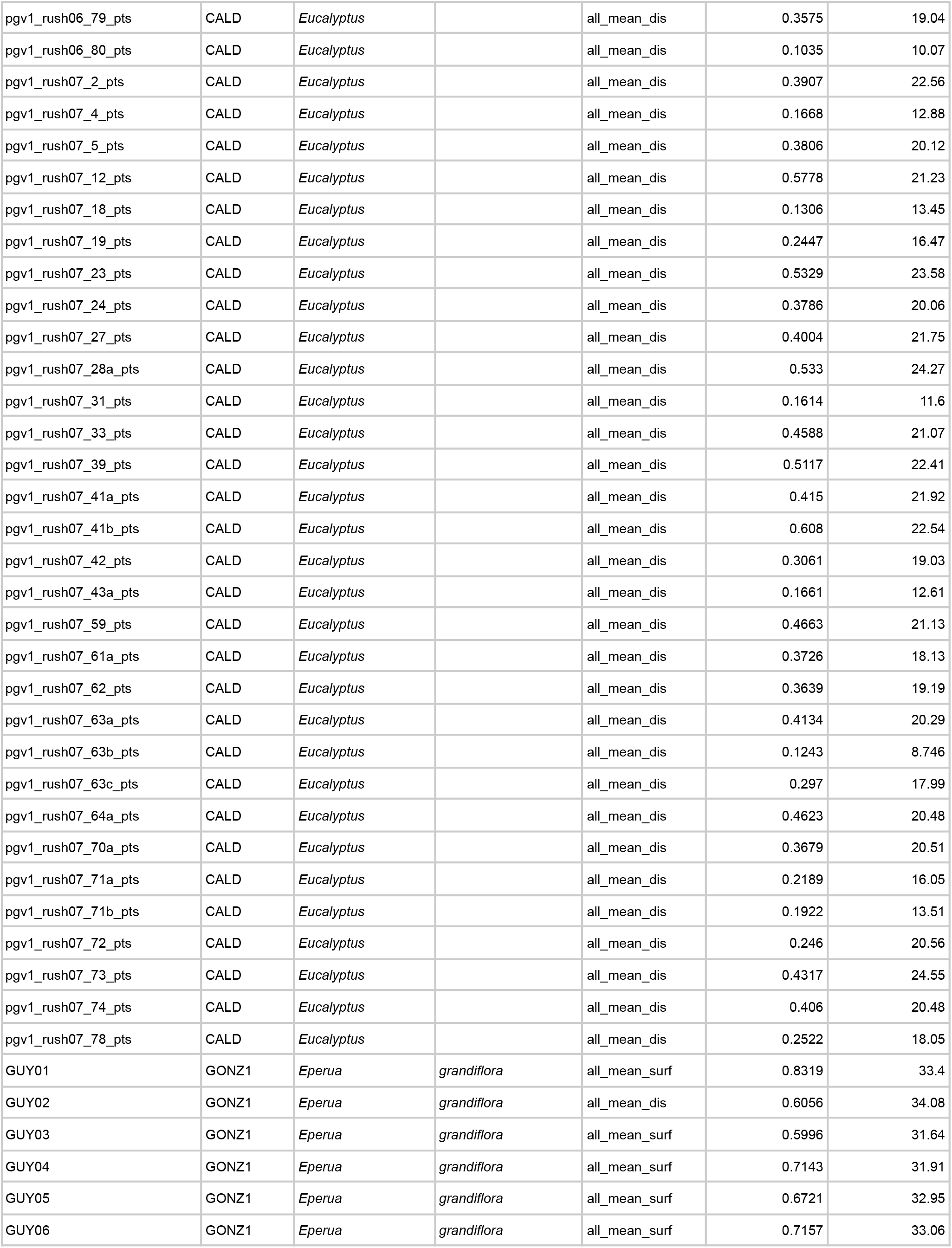
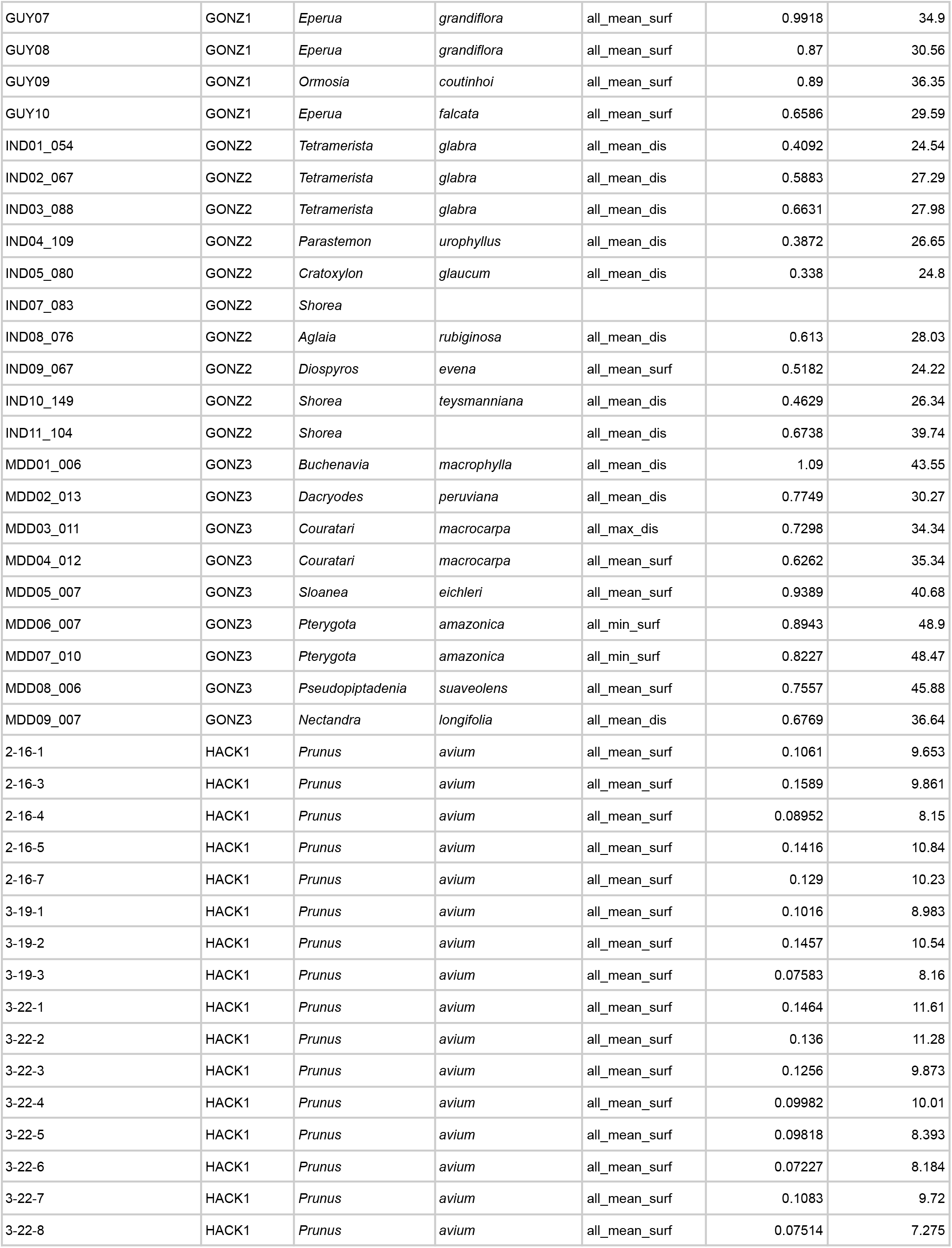
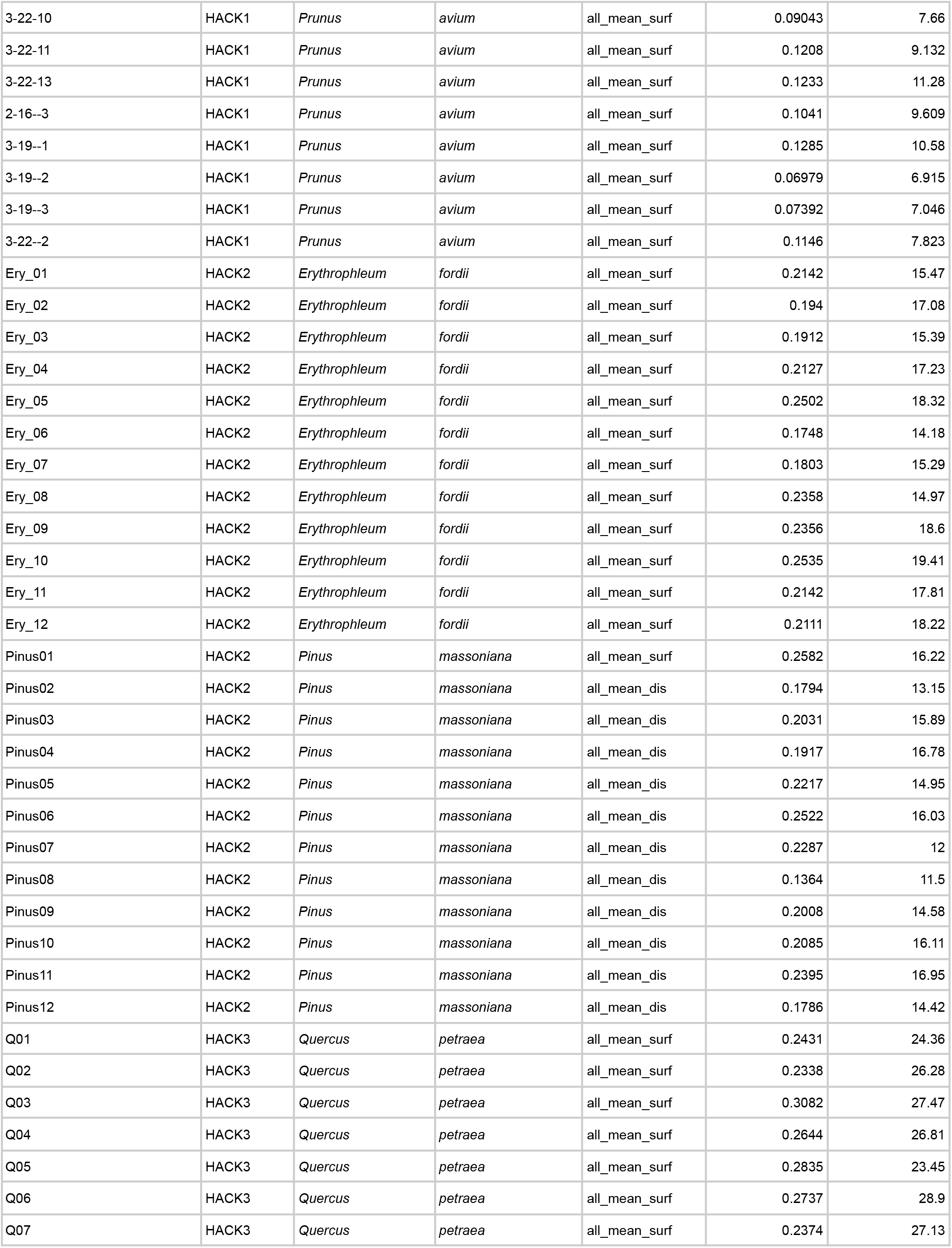
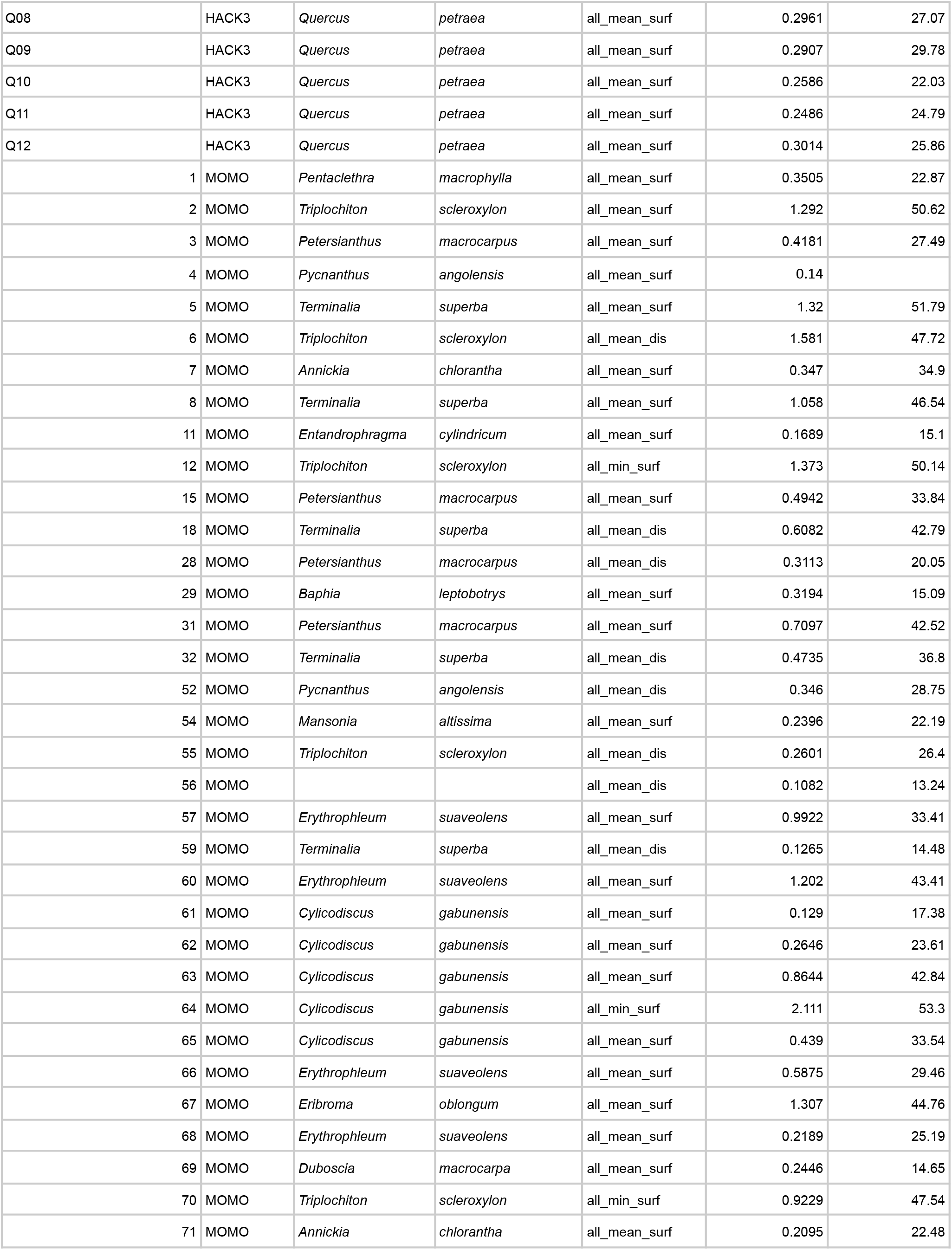
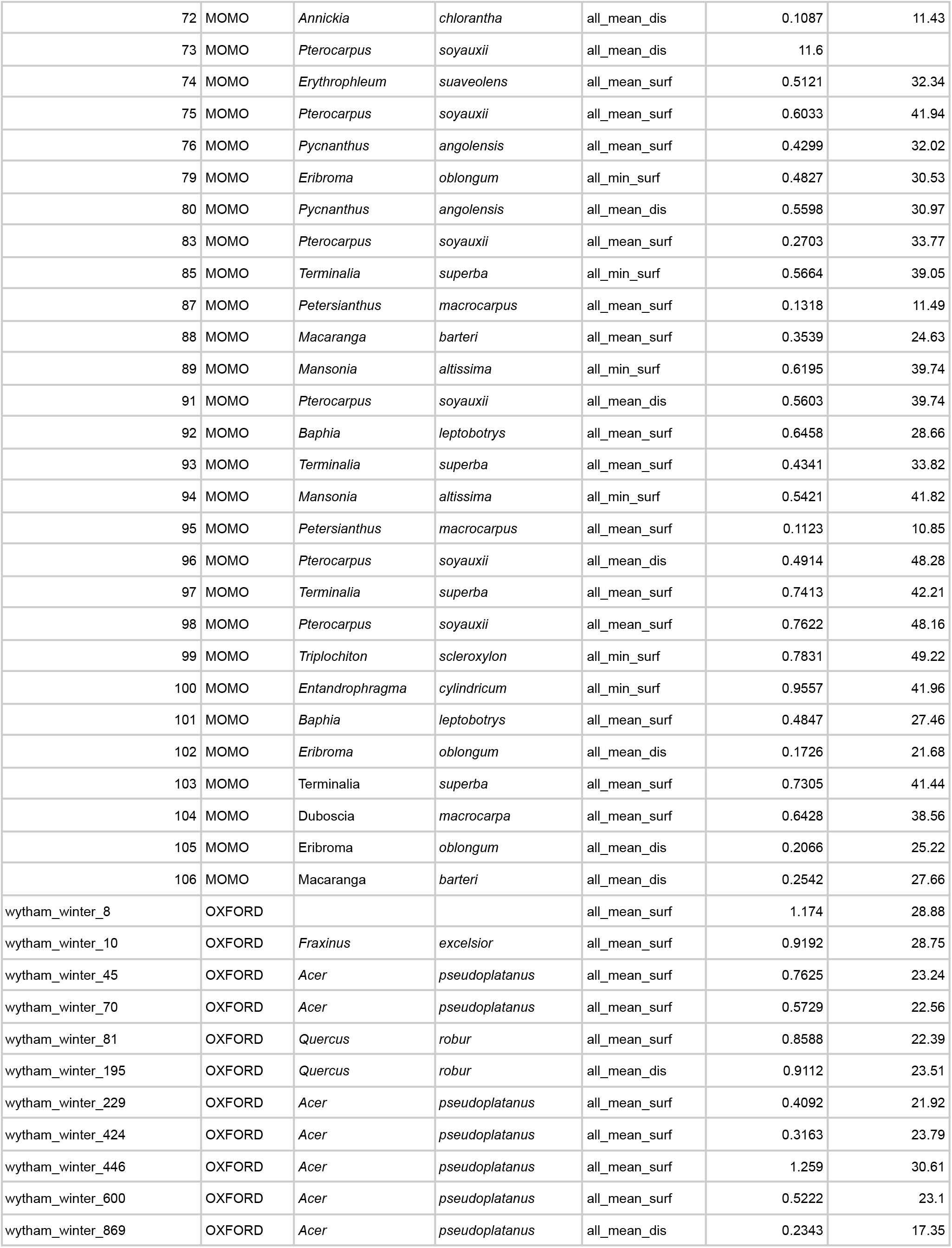
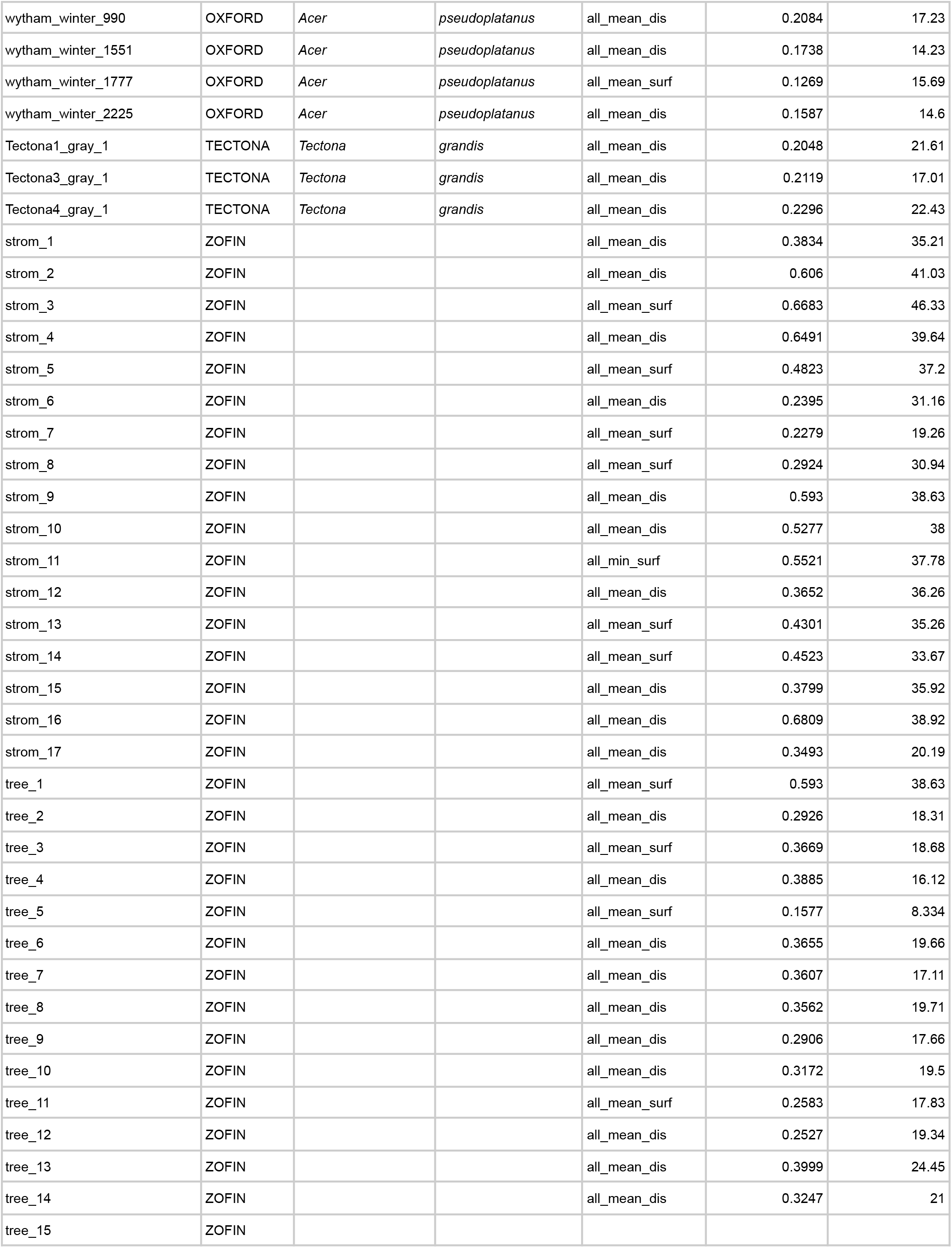
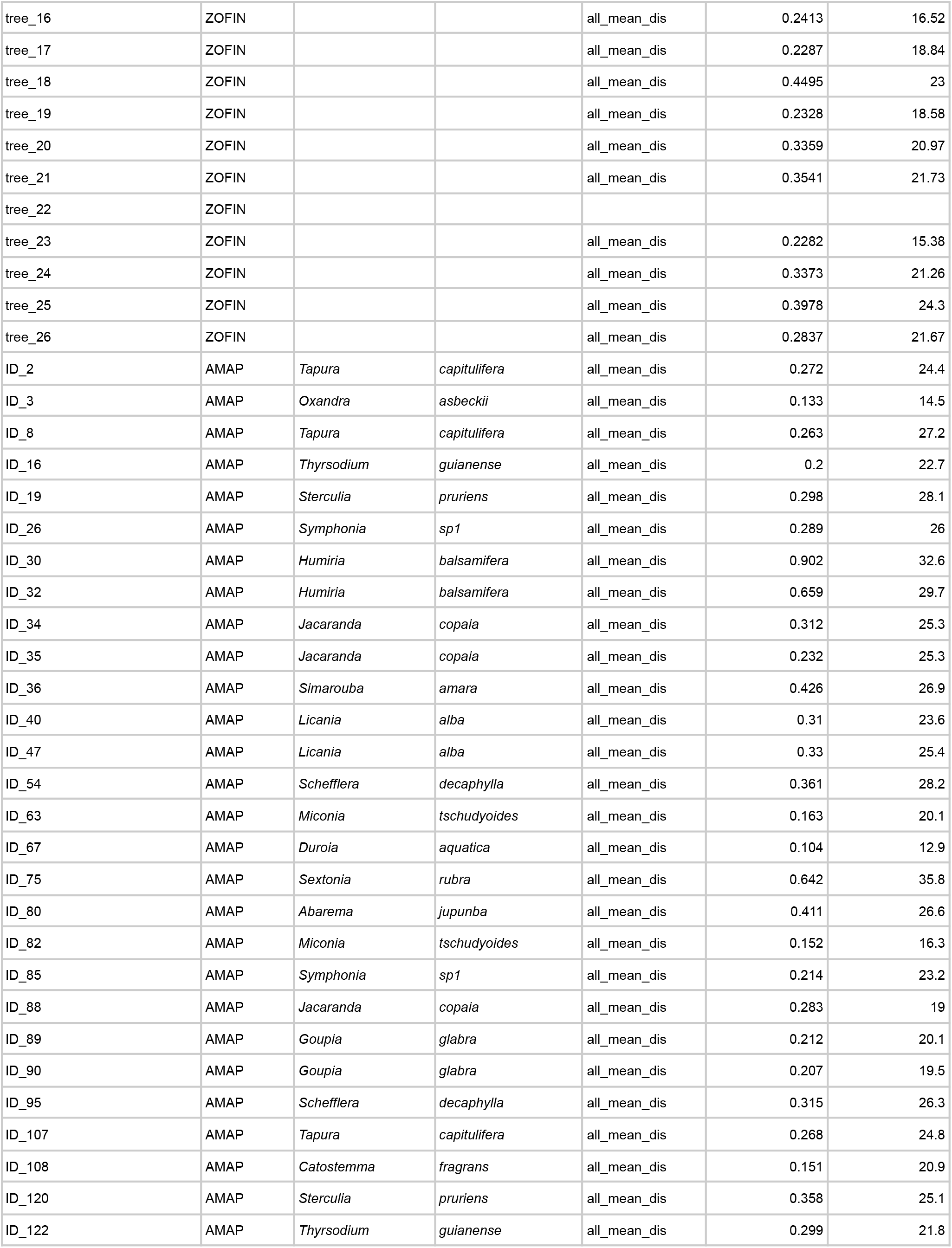
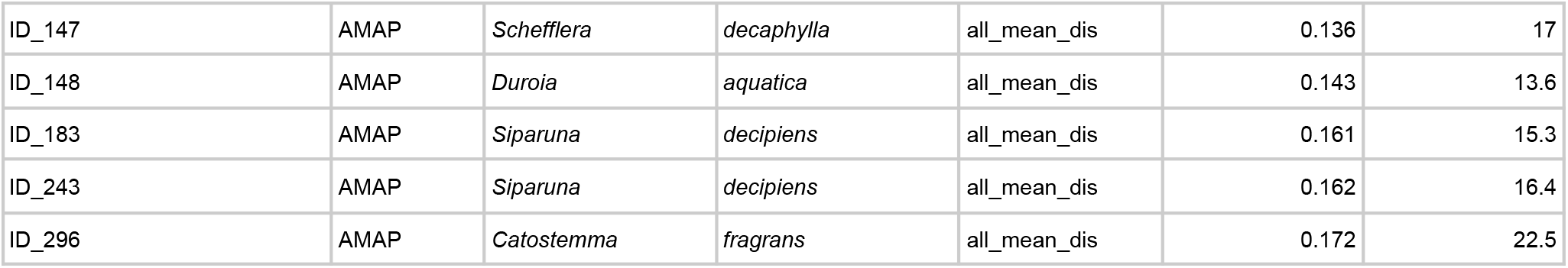
Individual tree data points with species information, TreeQSM optimization criterion, and DBH and height measured derived from QSM.

**Table S4:** List of hand-measured plant network data.

| Name | Species | Basal radius (m) | Source |
| --- | --- | --- | --- |
| GUY01 | *Eperua grandiflora* | 0.87 | Lau et al 2019 |
| GUY02 | *Eperua grandiflora* | 0.5 | Lau et al 2019 |
| GUY03 | *Eperua grandiflora* | 0.6 | Lau et al 2019 |
| GUY04 | *Eperua grandiflora* | 0.64 | Lau et al 2019 |
| GUY05 | *Eperua grandiflora* | 0.68 | Lau et al 2019 |
| GUY06 | *Eperua grandiflora* | 0.72 | Lau et al 2019 |
| GUY07 | *Eperua grandiflora* | 0.85 | Lau et al 2019 |
| GUY08 | *Eperua grandiflora* | 0.78 | Lau et al 2019 |
| GUY09 | *Ormosia coutinhoi* | 0.987 | Lau et al 2019 |
| GUY10 | *Eperua grandiflora* | 0.6 | Lau et al 2019 |
| balsa | *Ochroma pyramidale* | 0.375 | Bentley et al 2013 |
| df737g4 | *Pseudotsuga menziesii* | 0.00367 | Brummer et al. 2021 |
| df788g4 | *Pseudotsuga menziesii* | 0.0048 | Brummer et al. 2021 |
| df888g4 | *Pseudotsuga menziesii* | 0.0099 | Brummer et al. 2021 |
| df896g4 | *Pseudotsuga menziesii* | 0.00569 | Brummer et al. 2021 |
| dfg2 | *Pseudotsuga menziesii* | 0.00591 | Brummer et al. 2021 |
| maple1 | *Acer grandidentatum* | 0.00963 | von Allmen et al 2012 |
| maple2 | *Acer grandidentatum* | 0.00423 | von Allmen et al 2012 |
| maple3 | *Acer grandidentatum* | 0.00413 | von Allmen et al 2012 |
| maple4 | *Acer grandidentatum* | 0.01172 | von Allmen et al 2012 |
| maple5 | *Acer grandidentatum* | 0.00723 | von Allmen et al 2012 |
| oak1 | *Quercus gambelii* | 0.0041 | von Allmen et al 2012 |
| oak2 | *Quercus gambelii* | 0.00466 | von Allmen et al 2012 |
| oak3 | *Quercus gambelii* | 0.0056 | von Allmen et al 2012 |
| oak4 | *Quercus gambelii* | 0.00586 | von Allmen et al 2012 |
| oak5 | *Quercus gambelii* | 0.00835 | von Allmen et al 2012 |
| rob1212g5 | *Robinia neomexicana* | 0.00448 | Brummer et al. 2021 |
| rob1 | *Robinia neomexicana* | 0.00883 | Brummer et al. 2021 |
| rob2 | *Robinia neomexicana* | 0.00492 | Brummer et al. 2021 |
| rob3g5 | *Robinia neomexicana* | 0.00454 | Brummer et al. 2021 |
| rob4g5 | *Robinia neomexicana* | 0.00518 | Brummer et al. 2021 |
| wf1017g4 | *Abies concolor* | 0.00508 | Brummer et al. 2021 |
| wf256 | *Abies concolor* | 0.00562 | Brummer et al. 2021 |
| wf288g2 | *Abies concolor* | 0.00778 | Brummer et al. 2021 |
| wf327g2 | *Abies concolor* | 0.00731 | Brummer et al. 2021 |
| wf755g4 | *Abies concolor* | 0.00536 | Brummer et al. 2021 |
| wf991g4 | *Abies concolor* | 0.00583 | Brummer et al. 2021 |
| wp738g4 | *Pinus strobus* | 0.00934 | Brummer et al. 2021 |
| wp760g5 | *Pinus strobus* | 0.00517 | Brummer et al. 2021 |
| wp762 | *Pinus strobus* | 0.00778 | Brummer et al. 2021 |
| wp773 | *Pinus strobus* | 0.00872 | Brummer et al. 2021 |
| wp891g4 | *Pinus strobus* | 0.00725 | Brummer et al. 2021 |
| pinon | *Pinus edulis* | 0.114 | Bentley et al 2013 |
| pond03_edited | *Pinus ponderosa* | 0.03 | Bentley et al 2013 |
| pond05_edited | *Pinus ponderosa* | 0.043 | Bentley et al 2013 |
| pond06_edited | *Pinus ponderosa* | 0.048 | Bentley et al 2013 |
| pond07_edited | *Pinus ponderosa* | 0.062 | Bentley et al 2013 |
| pond16_edited | *Pinus ponderosa* | 0.028 | Bentley et al 2013 |

## Notes

### Competing Interest Statement

The authors have declared no competing interest.

